# Comparative metabolomics of released pollen during dispersal reveals metabolic adaptations to cold and heat stress

**DOI:** 10.64898/2025.12.31.697197

**Authors:** Jena Rutuparna, Johny Ijaq, Ashif Ali, Divya K Unnikrishnan, Irfan Ahmad Ghazi Raj

**Author notes:** Corresponding author: Jena Rutuparna. Co-corresponding author: Irfan Ahmad Ghazi.

## Abstract

Heat and cold stress can severely impair released pollen, reducing pollen viability and ultimately limiting fertilization and crop productivity. Following release, pollen is directly exposed to fluctuating environmental temperatures, necessitating adaptive metabolic mechanisms to sustain viability during dispersal. In response to temperature stress, pollen undergoes metabolic reprogramming that supports biochemical and physiological adaptation. However, the metabolic basis of pollen tolerance to extreme temperatures remains incompletely understood. In this study, biochemical assays combined with LC-MS-based untargeted metabolomic profiling were employed to investigate metabolic changes in released pollen exposed to low (15 °C) and high (35 °C) temperature stress. A total of 484 metabolites were detected, of which 147 were significantly altered between cold– and heat-stressed pollen, including 61 upregulated and 86 downregulated metabolites. Differentially regulated metabolites spanned multiple classes, including amino acids, flavonoids, long-chain fatty acids, sugars, and polyamines. Pathway enrichment analysis highlighted biologically relevant perturbations in amino acid and nucleic acid metabolism, including purine metabolism, arginine biosynthesis, and glutathione metabolism. Collectively, these findings demonstrate temperature-dependent metabolic adjustments in released pollen and provide new insights into the biochemical strategies underlying pollen tolerance to heat and cold stress. This work advances our understanding of pollen metabolic adaptation during dispersal and provides a foundation for identifying metabolic indicators relevant to crop fertility under changing climatic conditions. These findings provide a metabolomic framework for understanding pollen thermotolerance during dispersal and offer a resource for future studies on reproductive resilience under climate change.

## Introduction

Abiotic stresses such as extreme cold and heat pose major challenges to plant reproductive success and global agricultural productivity (Zhang et al., 2025). Among reproductive tissues, pollen viability and functionality are particularly sensitive to environmental fluctuations and often determine fertilization efficiency under stress conditions. Consequently, understanding pollen stress responses is critical for improving crop resilience and elucidating plant adaptation mechanisms (Chaturvedi et al., 2021). Wind-pollinated species, while widespread in various ecosystems, typically rely on effective pollen dispersal and survival to ensure reproductive success. *Putranjiva roxburghii* Wall., a dioecious member of the Euphorbiaceae family predominantly distributed in Asian tropical regions, represents a typical wind-pollinated species (Wurdack et al., 2004).

Pollen development involves a series of tightly regulated stages, including microspore formation, pollen maturation, pollination, pollen–stigma interaction, fertilization, and seed set (de Graaf and Dewitte, 2019). While pollen development within floral organs has been extensively studied, the metabolic characteristics of released pollen exposed to environmental stress during dispersal remain poorly understood. During this phase, extreme temperature conditions can impair cellular energy balance, disrupt metabolic homeostasis, and ultimately compromise reproductive capacity (Rieu et al., 2017). Plants exposed to such unfavourable conditions often activate thermotolerance mechanisms during early developmental stages to ensure survival (Hasanuzzaman et al., 2013), reflecting the evolutionary pressure to develop adaptive metabolic strategies that enhance stress resilience (Koza et al., 2022). Since heterotrophic pollen development depends on tightly coordinated metabolic processes, both heat and cold stress can perturb key pathways involving carbohydrates, amino acids, lipids, plant hormones, alkaloids, and flavonoids (Sharma and Nayyar, 2016; Raza et al., 2021; Duan et al., 2025). Accordingly, stress-induced alterations in specific metabolites and biomolecules may play decisive roles in regulating pollination efficiency and temperature stress adaptation (Zandalinas et al., 2022).

Recent metabolic investigations have demonstrated the critical roles of metabolites in pollen development and environmental stress responses (Miura and Furumoto, 2013; Paupiere et al., 2017; Carrera et al., 2021). Carbohydrate reserves, including sucrose and other soluble sugars, play essential roles in stabilizing cell membranes and maintaining turgor in pollen grains, and changes in pollen carbohydrate composition have been linked to desiccation tolerance and pollen viability during dispersal (Pacini et al., 2006). In rice, metabolomic analyses have shown that sugar insufficiency is a major factor contributing to reproductive failure in floral organs, including pollen grains, under drought and heat stress conditions (Rezaul et al., 2019; Lawas et al., 2019; Mo et al., 2023). Supporting these observations, untargeted metabolomic analysis of tomato pollen revealed that heat stress induces pronounced alterations in the pollen metabolome, particularly affecting flavonoids and other secondary metabolites associated with pollen viability and fertilization potential (Paupiere et al., 2017). Similarly, studies in maize have reported that heat stress exerts prolonged negative effects on pollen development and subsequent fertilization success (Smith et al., 2019). Moreover, low-temperature exposure has been reported to alter amino acid and carbohydrate metabolism in pollen, leading to impaired pollen germination and tube growth (Dai et al., 2022).

Beyond carbohydrates, several other metabolite classes have been implicated in pollen development and stress tolerance. In wind-pollinated species such as birch and maize, polyamines such as putrescine, spermidine, and spermine are abundant on the pollen surface, frequently occurring as conjugates with hydroxycinnamic acids (Liu et al., 2022). These compounds play essential roles in pollen development, cell wall formation, and sporopollenin deposition during pollen tube growth (Elejalde-Palmett et al., 2015; Liu et al., 2024). In addition to their developmental roles, polyamines have been shown to accumulate in plant tissues under diverse stress conditions, and modulation of polyamine metabolism has been associated with enhanced tolerance to heat, cold, and oxidative stresses (Alcazar et al., 2010). Earlier studies have also demonstrated that polyamines are closely linked to the phenylpropanoid pathway, facilitating exine biosynthesis through coordinated phenolic metabolism (Dong et al., 2021). Flavonols, a subgroup of flavonoids derived from the phenylpropanoid pathway, have been shown to maintain reactive oxygen species (ROS) homeostasis in pollen, thereby protecting pollen viability, germination, and tube growth under elevated temperature stress (Muhlemann et al., 2018). Despite these advances in understanding metabolite-mediated regulation of pollen development and stress responses, comparative metabolomic studies focusing specifically on released or shed pollen exposed to extreme temperatures remain limited (Paupiere et al., 2017; Wang et al., 2022).

So far, most studies examining pollen responses to temperature stress have focused on pollen grains developing within floral tissues, i.e., in a plant-dependent context (Rieu et al., 2017; Stokes and Geitmann, 2024). After maturation, however, pollen is thought to enter a developmental arrest phase that facilitates dispersal under diverse environmental conditions (Pacini and Dolferus, 2019; Williams and Brown, 2018). During this stage, desiccation tolerance mechanisms provide crucial adaptations that enable pollen survival during dispersal (Franchi et al., 2011; Pacini and Dolferus, 2019; Fan et al., 2025), suggesting the presence of intrinsic adaptive strategies that help maintain pollen viability outside the parental plant. Nevertheless, the molecular, biochemical, cellular, and metabolic adaptations that pollen may undergo during this developmental arrest phase, particularly under temperature stress, remain largely unexplored.

Despite advances in floral and pollen biomechanics, intraspecific variation in structural traits that promote effective pollen release and transport under stress remains underexplored (Timerman & Barrett, 2021). In this study, we aim to investigate both the structural changes that enable pollen dispersal during wind pollination and the metabolites and bioactive molecules that contribute to pollen function under cold and heat stress. Using untargeted LC–MS-based metabolite profiling of *P. roxburghii* pollen exposed to both cold and heat stress, complemented by quantitative assays for polyamines, hormones, and amino acids, we evaluate metabolic signatures and structural changes associated with stress tolerance. By focusing specifically on released pollen exposed to adverse environmental conditions, our work provides new insights into the molecular and biochemical strategies that support adaptive traits such as pollen resilience, functionality, and viability, which are crucial for successful wind pollination under fluctuating temperature conditions.

## Materials and methods

### 2.1 Pollen collection

Ornamentally grown *Putranjiva roxburghii* plants maintained at the University of Hyderabad, Hyderabad, India, were used in this study. Samples were collected in April 2019. The developmental stages of male flowers were monitored to determine the pollen release period, following the method described by Shivanna and Rangaswamy (2012). Mature pollen grains were collected from dehiscent anthers of fully bloomed flowers.

### 2.2 Experimental setup

Freshly collected pollen grains were transferred into 1.5 mL Eppendorf tubes. For stress treatments, two sets of tubes were incubated at approximately 15 °C (cold stress), and two additional sets were incubated at approximately 35 °C (heat stress) for 24 h. Following incubation, pollen samples were immediately processed for downstream experimental analyses.

### 2.3 Pollen viability assay

Pollen viability under cold and heat stress conditions was assessed following the method described by Muhlemann et al. (2018) with slight modifications. Briefly, pollen grains were resuspended in a solution containing 1.27 mM Ca(NO□)□, 290 mM sucrose, 0.16 mM boric acid, and 1 mM KNO (1% w/v), supplemented with 10 μL propidium iodide and 0.001% (w/v) fluorescein diacetate. Live and dead pollen were quantified using a flow cytometer (Muse™ Cell Analyzer; Merck Millipore).

### 2.4 Quantification of sugars, proline, hormones and polyamines

Total sugars were extracted following the method of Giannoccaro et al. (2006) and analyzed by HPLC (LC-20AD, Shimadzu, Japan). Separation was performed on an NH column (Shodex-Asahipak NH P-50-4E) using acetonitrile:water (70:30, v/v) as the mobile phase, at a flow rate of 0.8 mL/min. Sugars were detected using a photodiode array detector at 190 nm, and sucrose, fructose, and glucose were used as external standards.

Proline quantification was performed according to Guo and Trotter (2004). Extracted samples were derivatized with o-phthaladehyde (OPA) reagent (1:1:1) as described by McKerrow (2000) and analyzed by HPLC at a flow rate of 0.8 mL/min with UV detection at 262 nm.

For phytohormone analysis, collected pollen samples were ground in extraction buffer (2-propanol:H O:concentrated HCl, 2:1:0.002, v/v/v), followed by the addition of 1 mL dichloromethane. After centrifugation at 13,000 rpm for 5 min, 4 °C, the lower organic phase was collected, evaporated under nitrogen, and re-dissolved in methanol (Pan et al., 2010). Chromatographic separation was performed using solvent A (1% glacial acetic acid, v/v) and solvent B (100% acetonitrile) at a flow rate of 1.5 mL/min with an injection volume of 20 μL. The gradient program consisted of 0–55% acetonitrile over 25 min, followed by 100% acetonitrile for 3 min, and a final 1% acetic acid wash, with a total run time of 30 min. Standard phytohormones included indole-3-acetic acid, abscisic acid, gibberellic acids, salicylic acid, jasmonic acid, indole-3-butyric acid, 1-naphthaleneacetic acid, and kinetin (all from Sigma-Aldrich).

Polyamine analysis was performed by HPLC as described by Smith et al. (1985), using a 30 min linear gradient program. Quantification was based on the peak areas obtained from known standard concentrations.

### 2.5 Sample extraction for metabolome analysis

Approximately 1 mg of pollen was finely ground and extracted with 1 mL of extraction buffer consisting of methanol: chloroform: water (7:2:1, v/v/v). The mixture was vortexed for 30 min at 4 °C, and subsequently centrifuged at 13,000 rpm for 15 min at 4 °C. The resulting supernatant was transferred to a fresh tube and used for LC–MS analysis.

### 2.6 LC-MS analysis

Metabolite extracts derived from pollen samples exposed to heat and cold stress were analyzed using LC-MS. Chromatographic separation was performed using an Agilent 1290 Infinity II UHPLC system coupled with an Agilent 6520 quadrupole time-of-flight (Q-TOF) mass spectrometer (Agilent Technologies, USA). Separation was achieved using a ZORBAX RX-C18 reverse-phase HPLC column (5 µm; 4.6 mm × 150 mm; Agilent Technologies, USA). A 3 µL sample was injected at a flow rate of 0.2 mL/min. The mobile phases used were solvent A (0.1% formic acid in ddH2O) and solvent B (0.1% formic acid in acetonitrile). A linear gradient program was set as follows: 0 min, 5% B; 15 min, 15% B; 20 min, 35% B; 27 min, 50% B; 35 min, 70% B; 39 min, 85% B; 45 min, 15% B; and re-equilibration to 5% B at 60 min. Data were acquired in both positive and negative ionization modes in separate runs over a mass range of 50-1500 Da, with a scan rate of 0.9 spectra/sec. Collision energy was determined using a linear interpolation formula with a slope of 3 and an offset of 10 V. The electrospray ionization (ESI) capillary voltage was set to 3.5 kV. Dual ESI source parameters included a drying gas flow rate of 5.0 L/min, a nitrogen gas temperature of 325 °C, and the nebulizer gas pressure of 30 psig. MS-TOF parameters were set as follows: the fragmentor voltage of 175 V, the skimmer voltage of 65 V, and the Oct 1 RF Vpp of 750 V. The overall workflow for the metabolomics study is summarized in Figure 1.

**Figure 1.**
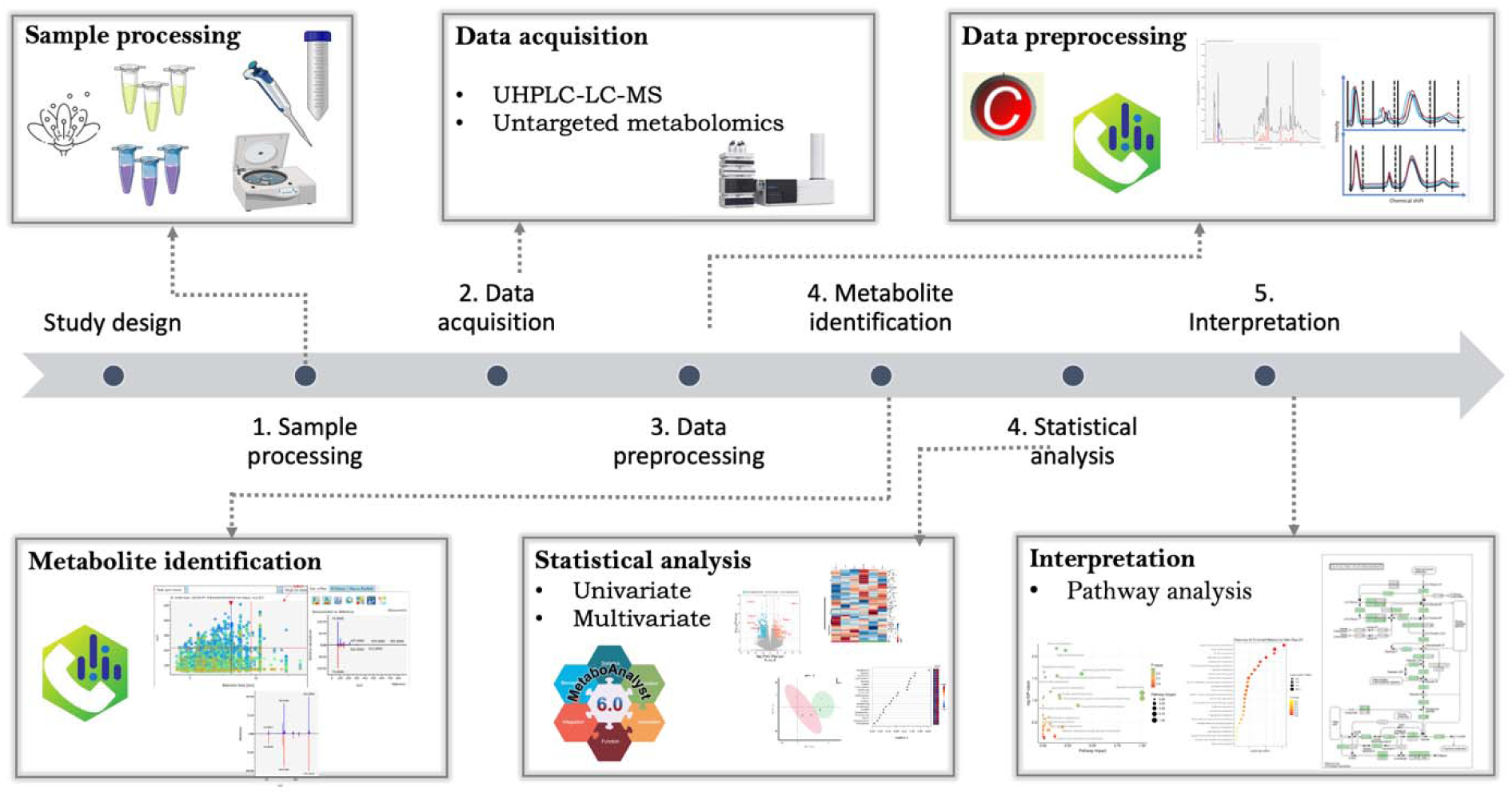
Experimental workflow for LC-MS-based metabolomics analysis of pollen under cold (R15) and heat (R35) stress.

### 2.7 Data processing and metabolite identification

Raw LC-MS data were pre-processed, including peak detection, deconvolution, and alignment, using MS DIAL (v 5.5.250404) software (Tsugawa et al., 2015). All the RAW data files were first converted to suitable file format (.abf) using the ABF converter (https://www.reifycs.com/abfconverter). Metabolite annotation was performed using a custom plant-specific and high-confidence MS/MS spectral libraries: RIKEN PlaSMA – Authentic Standards, which includes validated spectra of plant metabolites; the Fiehn/Vaniya Natural Product Library, which covers a broad range of plant secondary metabolites; and the ReSpect library, which contains spectra of phytochemicals such as flavonoids, phenolics, and alkaloids. Default MS-DIAL parameters were used, with mass accuracy thresholds of 0.01 Da for MS1 and 0.025 Da for MS2. Only peaks with a height above 1000 and a mass slice width > 0.1 Da were considered. Retention time and MS1 tolerances for alignment were set to 0.1 min and 0.015 Da, respectively. Positive ion adducts included [M+H]+, [M+Na]+, [M+NH4]+, and [M+H–H O]+, while negative ion adducts included [M–H]– and [M–H-H_2_O]-.

Alignment results were exported for further curation. Metabolites annotated with IUPAC-style names were excluded from the final list, due to limited biological relevance, lack of common chemical names, or unclear occurrence in plant metabolomes. Only metabolites with recognizable names were retained for interpretation. Following that, irrelevant and redundant compounds were manually removed. Subsequently, for metabolites identified at multiple retention times, the most representative feature was selected based on the highest average peak area across all samples, indicating the most consistent and abundant signal. The curated metabolite list constituted the final dataset used for downstream analyses.

### 2.8 Data analysis

Statistical analyses were performed using MetaboAnalyst 6.0 (Pang et al., 2024). Data were median-normalized, log transformed, and auto-scaled prior to analysis. Differentially abundant metabolites were identified using a Student’s unpaired two-tailed t-test combined with fold-change analysis. Metabolites with a fold change ≥ 2.0 and p-value < 0.05 were considered significant. Principal component analysis (PCA) was performed to visualize the overall clustering pattern of the samples. Hierarchical clustering heatmap was used to assess variance among samples. Pathway analysis was performed to identify significantly affected metabolic pathways.

### 2.9 Multivariate statistical analysis

Supervised multivariate analysis was performed using partial least squares discriminant analysis (PLS-DA) as an exploratory approach to identify metabolites that contributed most to the separation between cold– and heat-stressed pollen sample groups. This approach allowed to validate the robustness of the identified differentially expressed metabolites with the highest discriminatory power between the two groups. The model was assessed using cross-validation, and Variable Importance in Projection (VIP) scores were used to rank metabolites contributing to group separation

To further prioritize metabolites for biological interpretation, sparse PLS-DA (sPLS-DA) was applied in an exploratory manner. This method introduces sparsity in the model, selecting only the most relevant metabolites while reducing noise. Two components with 20 features each were selected, and metabolites with loading threshold values > 0.2 were considered for downstream interpretation.

### 2.10 Statistical analysis

All experiments were conducted in biological triplicates. Statistical significance was determined using Student’s unpaired two-tailed t-test, with p-values < 0.05 considered significant. Spearman’s correlation analysis was used to evaluate relationships between sample groups. For PLS-DA, the optimal number of components was determined using five-fold cross-validation. Figures were generated using ggplot2 and EasyPubPlot (Tien et al., 2025).

## Results

### 3.1 Effect of temperature on pollen viability

Pollen viability was assessed under two temperature conditions: cold stress (15 °C; R15) and heat stress (35 °C; R35). Reactive oxygen species (ROS) activity was examined to evaluate temperature-induced cellular damage, and pollen viability was quantified using fluorescence-activated cell sorting (FACS). Pollen viability was determined using propidium iodide (PI) and fluorescein diacetate (FDA) staining, where the absence of esterase activity and positive PI staining indicated non-viable pollen grains.

Released pollen exposed to cold stress (R15) exhibited a viability of 71%, whereas pollen subjected to heat stress (R35) showed a reduced viability of 55% (Figure 2). Overall, pollen viability decreased under both cold and heat stress conditions, with a more pronounced reduction observed under heat stress.

**Figure 2.**
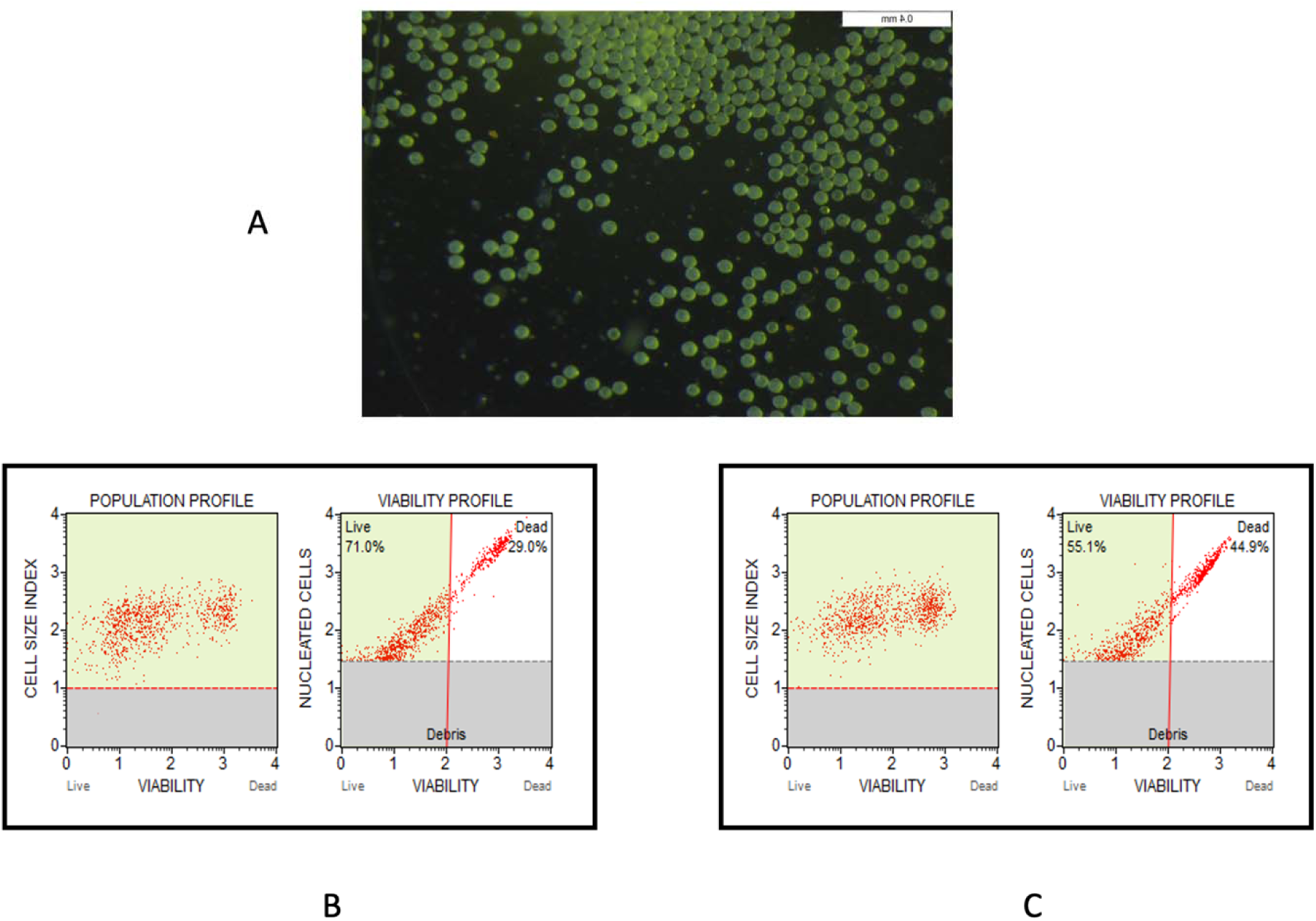
Pollen viability analysis under different temperature conditions using flow cytometry (FACS). (A) Pollen grains of *Putranjiva roxburghii*. (B) Flow cytometric analysis of pollen viability under cold stress (R15). (C) Flow cytometric analysis of pollen viability under heat stress (R35).

### 3.2 Sucrose accumulation is higher in heat-stressed pollen

Glucose, fructose, and sucrose were detected in pollen under both temperature conditions. Among these carbohydrates, sucrose was the most abundant sugar. Glucose levels under cold stress (R15) were 9.71 mg g□¹, whereas heat-stressed pollen (R35) showed a lower glucose content of 6.78 mg g□¹ (Supplementary Figure S1).

Sucrose accumulation increased under heat stress, with concentrations of 10.13 mg g□¹ at R15 and 10.93 mg g□¹ at R35 (Supplementary Figure S1). Fructose content was comparatively low, measuring 0.53 mg g□¹ under cold stress and 0.23 mg g□¹ under heat stress (Supplementary Figure S1).

### 3.3 Higher accumulation of proline in cold-stressed pollen

Proline content decreased under heat stress relative to cold stress. Released pollen under cold stress (R15) contained 5.65 nmol mL□¹ of proline, whereas heat-stressed pollen (R35) exhibited a lower concentration of 2.14 nmol mL□¹ (Supplementary Figure S2).

### 3.4 Plant hormone responses to cold and heat stress

Eight phytohormones including abscisic acid (ABA), indole-3-acetic acid (IAA), 1-naphthaleneacetic acid (NAA), indole-3-butyric acid (IBA), salicylic acid (SA), kinetin, gibberellic acid (GA□), and methyl jasmonate were quantified to assess hormonal responses in pollen under temperature stress. All eight hormones showed measurable changes following cold and heat treatments.

Among the analyzed hormones, GA□, kinetin, and SA exhibited comparatively higher concentrations across temperature conditions (Figure 3). IAA levels were 0.476 nmol g□¹ under cold stress (R15) and 0.722 nmol g□¹ under heat stress (R35) (Figure 3). IBA concentrations were 0.005 nmol g□¹ at R15 and 0.015 nmol g□¹ at R35 (Figure 4b). GA was the most abundant hormone, with concentrations of 479.86 nmol g□¹ under cold stress and 358.20 nmol g□¹ under heat stress (Figure 3). ABA levels were 0.028 nmol g□¹ at R15 and 0.047 nmol g□¹ at R35 (Figure 3). Methyl jasmonate and NAA showed minimal variation between cold and heat stress conditions (Figure 3).

**Figure 3.**
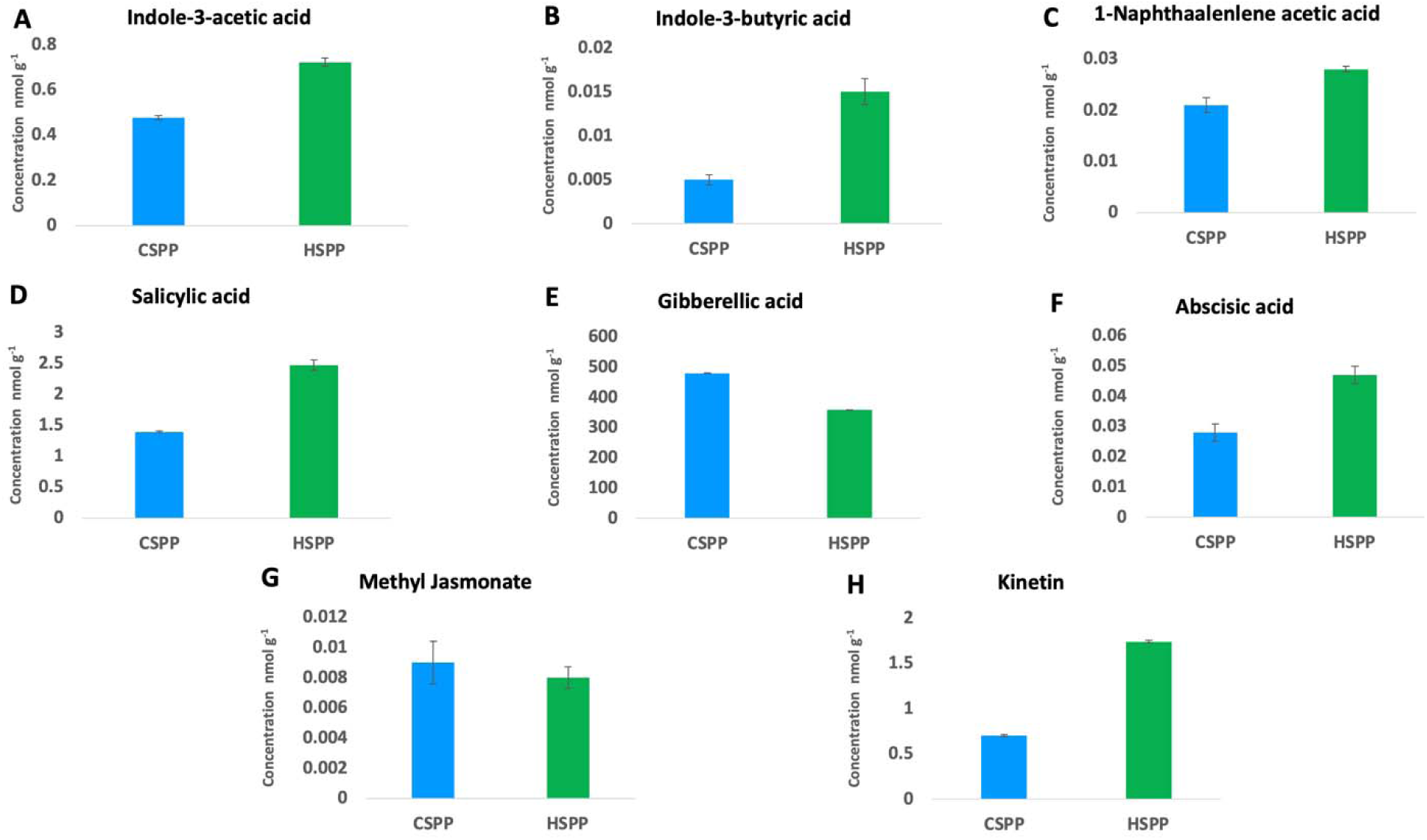
Quantification of phytohormones in pollen under cold (R15) and heat (R35) stress. Data represent mean ± SD (n = 3). (A) Indole-3-acetic acid (B) Indole-3-butyric acid (C) 1-Napthalene acetic acid (D) Salicylic acid (E) Gibberellic acid (F) Abscisic acid (G) Methyl jasmonate (H) Kinetin.

**Figure 4.**
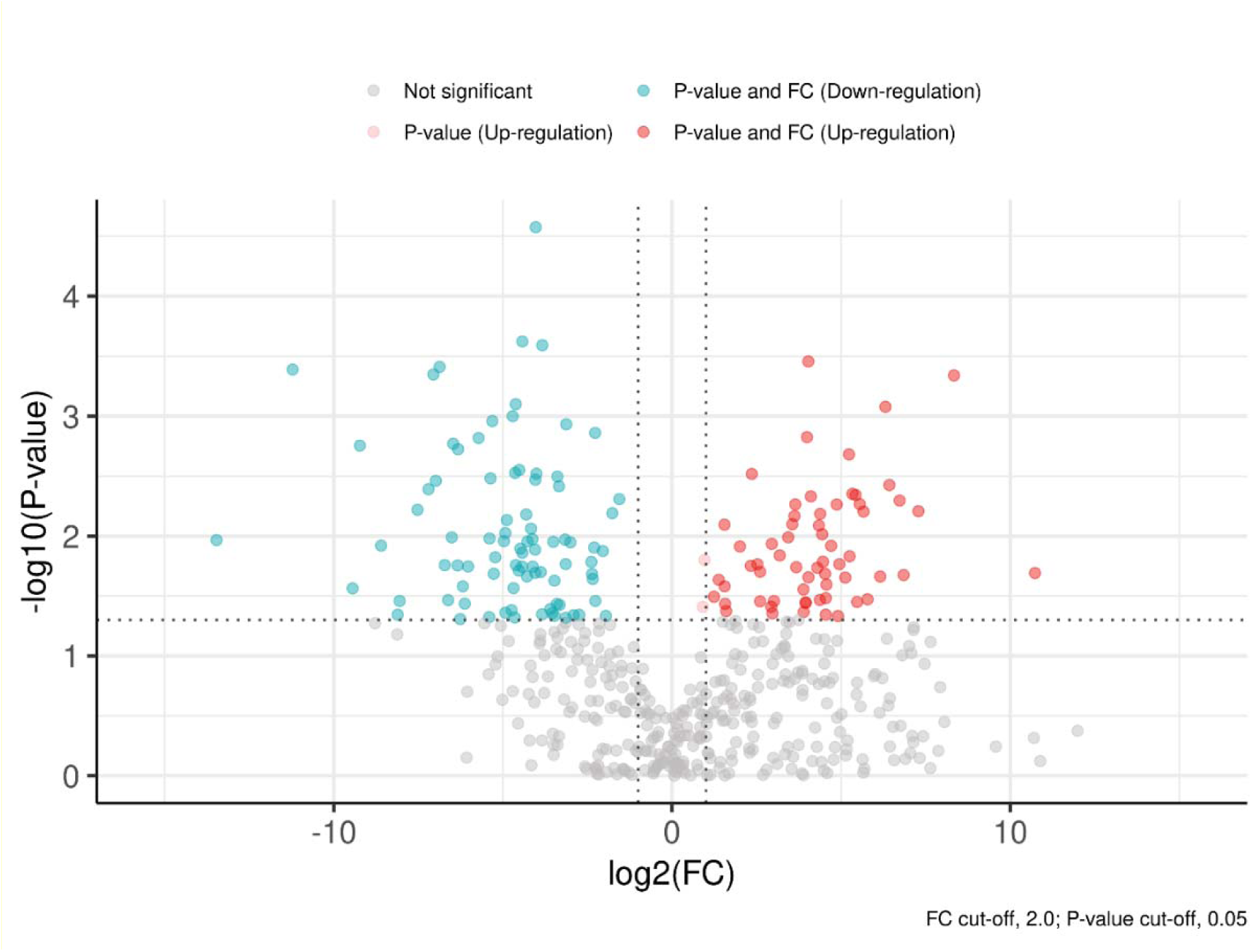
Volcano plot showing log fold change versus –log adjusted p-value, highlighting significantly altered metabolites.

### 3.5 Polyamine levels under cold and heat stress

Four major polyamines, including putrescine, spermidine, spermine, and cadaverine were quantified in released pollen under both temperature conditions. Putrescine was the most abundant polyamine, with concentrations of 178 nmol g□¹ under cold stress and 148 nmol g□¹ under heat stress (Supplementary Figure S3). Spermine was the second most abundant polyamine, measuring 56 nmol g□¹ at R15 and 44.6 nmol g□¹ at R35 (Supplementary Figure S3).

Cadaverine levels decreased under heat stress, from 1.06 nmol g□¹ at R15 to 0.488 nmol g□¹ at R35 (Supplementary Figure S3). Spermidine concentrations were 5.7 nmol g□¹ under cold stress and 7.62 nmol g□¹ under heat stress (Supplementary Figure S3), representing the lowest abundance among the analyzed polyamines.

### 3.6 Metabolomic profiling reveals temperature-dependent metabolic shifts

Untargeted metabolomic analysis of pollen subjected to cold and heat stress identified a total of 484 metabolites, including 478 detected in positive ion mode and 7 in negative ion mode. The major metabolite classes included amino acids and their derivatives, flavonoids and related compounds, and long-chain fatty acids (Supplementary Data S1).

Principal component analysis (PCA) was performed using all detected metabolites to visualize global metabolic patterns. PCA revealed distinct clustering between cold– and heat-stressed pollen sample groups, indicating consistent temperature-dependent shifts in the metabolome (Supplementary Figure S4). Heat-stressed samples clustered closely, whereas one cold-stressed replicate showed a slight deviation. The first two principal components explained 75.6% of the total variance (PC1: 49.0%; PC2: 26.6%) (Supplementary Data S2). Hierarchical clustering of the top 50 significantly altered metabolites further demonstrated clear group-level differences in metabolite abundance (Supplementary Figure S5).

Differential expression analysis identified 147 significantly altered metabolites (fold change >2.0, p < 0.05) between cold– and heat-stressed pollen samples, of which 61 were upregulated, and 86 were downregulated under cold stress (Supplementary Data S2). Notably, berberine (FC = 1701.3, *p* = 0.020), palmitic acid (FC = 323.4, *p* = 0.0004), and glycocholic acid (FC = 155.8, *p* = 0.006) were among the most upregulated metabolites under cold stress. In contrast, isopalmitic acid (FC = 8.8 × 10□□, *p* = 0.0108), karanjin (FC = 0.00042, *p* = 0.0004), and alizarin (FC = 0.0017, *p* = 0.0018) were strongly downregulated. These results are visualized in a volcano plot (Figure 4), and normalized abundance levels for representative metabolites are shown in box plots (Figures 5A and 5B). The top 20 upregulated and downregulated metabolites are listed in Table I.

**Figure 5.**
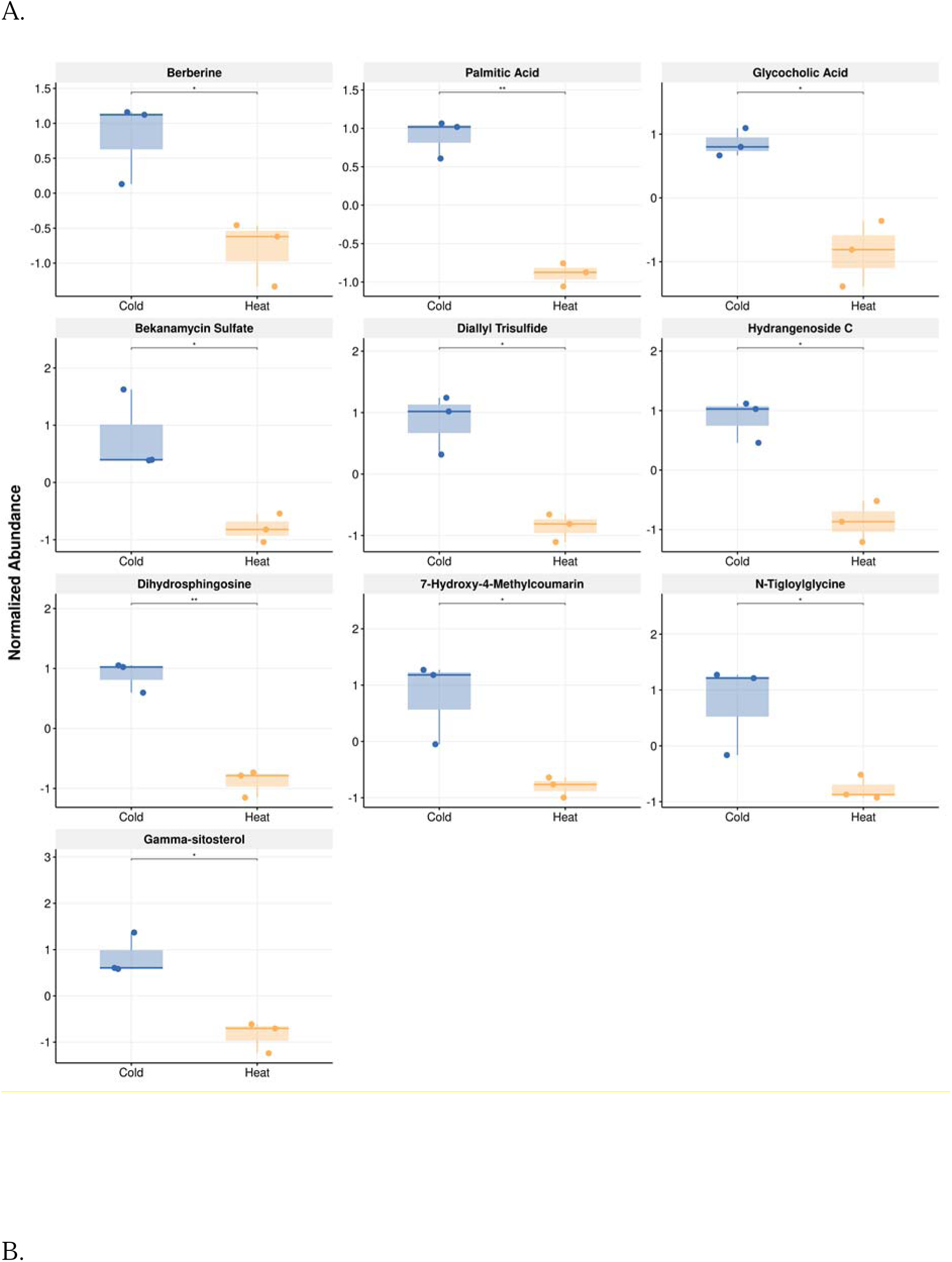
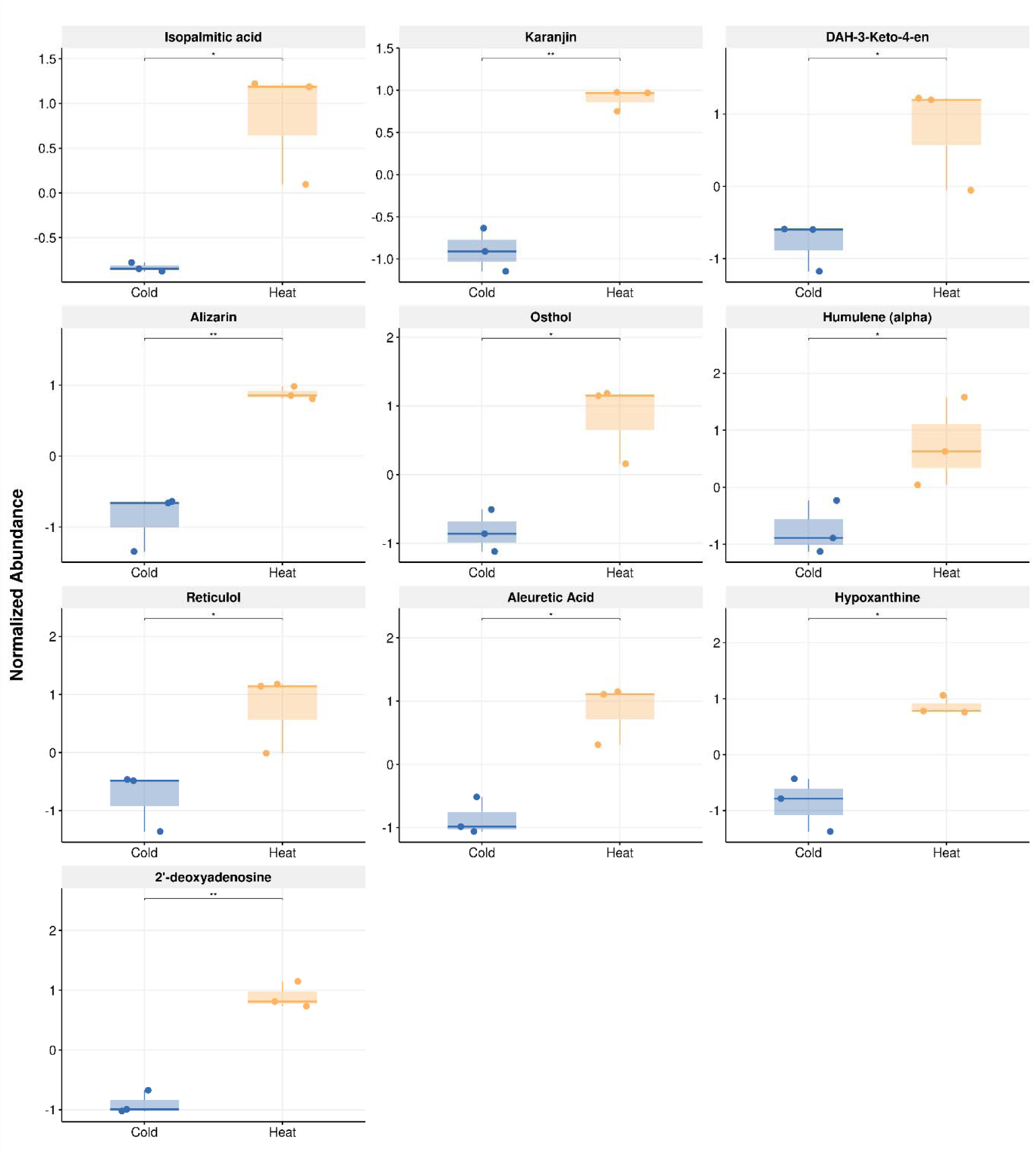
Box plots of the top 10 upregulated (5A) and downregulated metabolites (5B), showing group-wise distribution and variability.

**Table 1.** List of top 20 upregulated and downregulated metabolites in cold stressed-pollen compared to heatstressed-pollen, ranked by fold change and log2FC.

| Metabolite | Regulation | Fold change | Log2(FC) | p-value |
| --- | --- | --- | --- | --- |
| Berberine | Up | 1701.3 | 10.732 | 0.020369 |
| Palmitic Acid | Up | 323.43 | 8.3373 | 0.00045736 |
| Glycocholic Acid | Up | 155.86 | 7.2841 | 0.0062019 |
| Bekanamycin Sulfate | Up | 115.35 | 6.8499 | 0.021151 |
| Diallyl Trisulfide | Up | 106.23 | 6.731 | 0.0050518 |
| Hydrangenoside C | Up | 86.081 | 6.4276 | 0.0037544 |
| Dihydrosphingosine | Up | 79.382 | 6.3107 | 0.00083594 |
| 7-Hydroxy-4-Methylcoumarin | Up | 71.319 | 6.1562 | 0.021757 |
| N-Tigloylglycine | Up | 55.039 | 5.7824 | 0.033749 |
| Gamma-sitosterol | Up | 50.655 | 5.6626 | 0.0062546 |
| Auranticin A | Up | 46.906 | 5.5517 | 0.0054111 |
| Inosine-5'-monophosphate | Up | 44.378 | 5.4718 | 0.035472 |
| Ormosanine | Up | 43.094 | 5.4294 | 0.0045449 |
| Osajin | Up | 40.355 | 5.3347 | 0.0044495 |
| Avocadyne 1-Acetate | Up | 38.068 | 5.2505 | 0.014765 |
| 7,8-Dihydromethysticin | Up | 37.618 | 5.2333 | 0.002085 |
| N-Glycylglycine | Up | 34.889 | 5.1247 | 0.0222 |
| Difucol Hexamethyl Ether | Up | 30.901 | 4.9496 | 0.017161 |
| L-Kynurenine | Up | 30.043 | 4.909 | 0.046536 |
| Betulin | Up | 29.247 | 4.8702 | 0.0054508 |
| Isopalmitic acid | Down | 8.84E-05 | -13.465 | 0.010805 |
| Karanjin | Down | 0.00042059 | -11.215 | 0.00040873 |
| DAH-3-Keto-4-en | Down | 0.0014384 | -9.4413 | 0.027321 |
| Alizarin | Down | 0.0016676 | -9.228 | 0.001762 |
| Osthol | Down | 0.0025744 | -8.6015 | 0.012024 |
| Humulene (alpha) | Down | 0.0036208 | -8.1095 | 0.045376 |
| Reticulol | Down | 0.0037673 | -8.0522 | 0.034801 |
| Aleuretic Acid | Down | 0.0054304 | -7.5247 | 0.0060453 |
| Hypoxanthine | Down | 0.0068013 | -7.2 | 0.0040644 |
| 2'-deoxyadenosine | Down | 0.0075256 | -7.054 | 0.00044906 |
| Citrulline | Down | 0.0079064 | -6.9828 | 0.0034584 |
| Biliverdin | Down | 0.0085419 | -6.8712 | 0.00038834 |
| Agelasine | Down | 0.0094985 | -6.7181 | 0.017484 |
| Deferrioxamine E | Down | 0.010182 | -6.6179 | 0.03426 |
| Estrone | Down | 0.010961 | -6.5115 | 0.010259 |
| N-Formyl-L-Methionine | Down | 0.011279 | -6.4702 | 0.0016989 |
| Carnosine | Down | 0.012361 | -6.3381 | 0.01757 |
| Anabasamine Hydrochloride | Down | 0.012472 | -6.3252 | 0.0018843 |
| Isopentenyladenine | Down | 0.013027 | -6.2623 | 0.049291 |
| Glutamine (L) | Down | 0.013745 | -6.1849 | 0.026285 |

Exploratory supervised analysis using partial least squares-discriminant analysis (PLS-DA) showed clear separation between cold– and heat-stressed pollen samples, indicating distinct metabolic alterations induced by temperature stress (Supplementary Figure S6). The first two components explained 74.2% of the variance (Component 1 = 47.2%, Component 2 = 27.0%). Metabolites with Variable Importance in Projection (VIP) scores ≥ 1.2 were considered contributors to group separation. This threshold identified 158 metabolites (32% of total features) that potentially contribute to the pollen’s metabolic response to temperature stress. Among these, caryophyllene (VIP = 1.510), piperidine (VIP = 1.497), theobromine (VIP = 1.496), cupressuflavone (VIP = 1.493) and biliverdin (VIP = 1.492) were ranked among the most discriminative features (Supplementary Figure S6, Supplementary Data S2). Model performance was evaluated using cross-validation, yielding R^2^ and Q^2^ values of 0.99 and 0.66, respectively (Supplementary Figure S6). Of the 147 differentially expressed metabolites, 145 (excluding Avocadyne 1-Acetate and Anisatin), also exhibited VIP scores greater than 1.2, indicating strong concordance between univariate and multivariate analyses (Supplementary Data S2).

Sparse PLS-DA (sPLS-DA) further identified 20 key metabolites from Component 1 and 20 from Component 2. Component 1 accounted for the majority of variance (43.5%) between temperature conditions and served as the primary axis of separation, whereas Component 2 captured residual variation (Supplementary Data S2). Ten metabolites from Component 1 overlapped with differentially expressed metabolites and exhibited VIP scores ≥ 1.2, forming a high-confidence subset consistently identified across univariate and multivariate analyses (Table 2). Loadings of top ten metabolites contributing to group separation are shown in Supplementary Figure S7.

**Table 2.** Final panel of discriminatory metabolites selected by combining differential expression analysis (FC ≥ 2.0, p < 0.05), PLS-DA (VIP ≥ 1.2), and sparse PLS-DA (Component 1). The table shows fold change (FC), log2 fold change, p-values (t-test), VIP scores from PLS-DA, sPLS-DA loadings with group contribution. Negative sPLS-DA loadings are associated with cold stress, while positive loadings correspond to heat stress.

| Metabolite | Fold Change | Log2FC | p-value | VIP Score | sPLS-DA Loading (Contribution) |
| --- | --- | --- | --- | --- | --- |
| Palmitic Acid | 323.43 | 8.33 | 0.000457 | 1.5103 | -0.2682 (Cold stress) |
| Dihydrosphingosine | 79.38 | 6.31 | 0.000835 | 1.4808 | -0.20455 (Cold stress) |
| Cupressuflavone | 16.34 | 4.03 | 0.000349 | 1.4935 | -0.29089 (Cold stress) |
| Karanjin | 0.0004 | -11.2 | 0.000408 | 1.4916 | 0.27807 (Heat stress) |
| 2'-deoxyadenosine | 0.0075 | -7.05 | 0.000449 | 1.4904 | 0.26984 (Heat stress) |
| Biliverdin | 0.0085 | -6.87 | 0.000388 | 1.4923 | 0.28238 (Heat stress) |
| 3',4',5,7-tetrahydroxyflavone | 0.04063 | -4.62 | 0.000794 | 1.4817 | 0.21073 (Heat stress) |
| Piperidine | 0.04654 | -4.42 | 0.000238 | 1.4976 | 0.31843 (Heat stress) |
| Caryophyllene | 0.06129 | -4.02 | 0.000026 | 1.5103 | 0.40493 (Heat stress) |
| Theobromine | 0.07004 | -3.83 | 0.000256 | 1.4969 | 0.31357 (Heat stress) |

Pathway analysis was performed using the 145 differentially expressed metabolites and with VIP scores >1.2. Although none of the pathways met the false discovery rate (FDR) threshold of 0.05, several biologically relevant pathways exhibited low raw p-values, suggesting potential pathway-level perturbations (Figure 6). Notably, purine metabolism (p = 0.0066), arginine biosynthesis (p = 0.0073), and glutathione metabolism (p = 0.0227) emerged as top hits. These pathways suggest potential metabolic adaptations to temperature stress and warrant further investigation. Top ten enriched pathways are shown in Table 3. A complete list of enriched pathways is provided in Supplementary Data S2.

**Figure 6.**
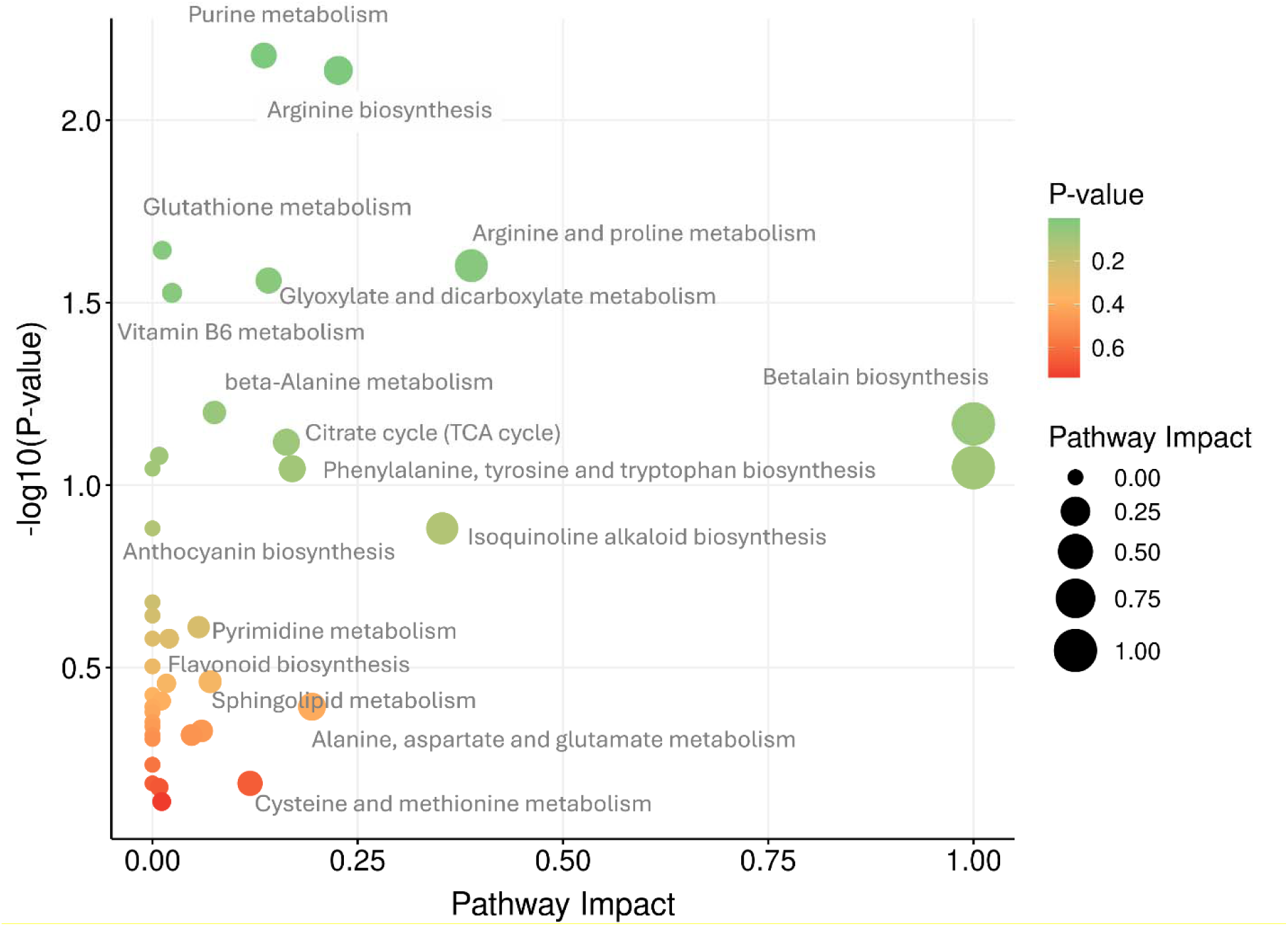
Pathway analysis of differentially expressed metabolites. The x-axis represents pathway impact based on topology analysis, and the y-axis shows –log10(p-value) from enrichment analysis. Circle size indicates pathway impact, while color intensity reflects statistical significance (This figure can be pushed to supplementary data)

**Table 3.** Pathway analysis results from MetaboAnalyst using KEGG database. Top 10 enriched metabolic pathways were listed. Pathways are ranked by p-value and impact score from topology analysis.

| **Description** | **Hits** | **Total** | **Impact** | **Raw p** | **FDR** |
| --- | --- | --- | --- | --- | --- |
| Purine metabolism | 6 | 76 | 0.13562 | 0.0066512 | 0.33978 |
| Arginine biosynthesis | 3 | 18 | 0.22633 | 0.007307 | 0.33978 |
| Glutathione metabolism | 3 | 27 | 0.01195 | 0.022717 | 0.46073 |
| Arginine and proline metabolism | 3 | 28 | 0.38838 | 0.025045 | 0.46073 |
| Glyoxylate and dicarboxylate metabolism | 3 | 29 | 0.14137 | 0.027498 | 0.46073 |
| Vitamin B6 metabolism | 2 | 12 | 0.02381 | 0.029724 | 0.46073 |
| beta-Alanine metabolism | 2 | 18 | 0.0754 | 0.063202 | 0.6447 |
| Betalain biosynthesis | 1 | 3 | 1 | 0.067931 | 0.6447 |
| Citrate cycle (TCA cycle) | 2 | 20 | 0.16309 | 0.076271 | 0.6447 |
| Zeatin biosynthesis | 2 | 21 | 0.00813 | 0.083104 | 0.6447 |

## Discussion

Climate change has led to significant shifts in environmental temperatures, increasing the exposure of pollen grains to both heat and cold stress. These stresses can reduce pollen viability and germination, leading to poor fertilization and lower crop yields (Chaturvedi et al., 2021). While plant-level responses to temperature stress have been extensively studied, comparatively little is known about the metabolic adaptations of pollen during wind pollination (Rosbakh et al., 2018; Pacini and Dolferus, 2019). During wind pollination, pollen in flight experiences rapid and extreme temperature variations, which can be highly stressful, challenging pollen viability and reproductive success (Muhlemann et al., 2018; Pacini and Dolferus 2019). To withstand these conditions and ensure successful fertilization, pollen must develop certain intrinsic metabolic and physiological adaptations (Zinn et al., 2010; Paupiere et al., 2014). In this context, the present study investigates these adaptive responses in pollen subjected to temperature stress, with a focus on released pollen, emphasizing the biochemical and metabolic mechanisms that support pollination under changing environmental conditions. By integrating targeted biochemical analyses of sugars, phytohormones, and polyamines with untargeted metabolomic profiling, we examined cold-stressed (R15) and heat-stressed (R35) pollen to capture coordinated changes in both primary and secondary metabolism. These analyses revealed temperature-dependent biochemical and metabolic changes that appear to support pollen survival and function under fluctuating temperature conditions, highlighting key metabolites and pathways contributing to pollen thermotolerance.

Pollen viability assays revealed significantly higher viability in cold-stressed pollen (R15; 71%) compared with heat-stressed pollen (R35; 55%), indicating a stronger detrimental effect of heat stress. Such temperature-dependent reductions in pollen viability are consistent with previous reports in crop species including wheat, sorghum, and rice, where elevated temperatures impair pollen function and fertilization efficiency (Sarkar et al., 2021; Fahad et al., 2018; Djanaguiraman et al., 2018). These observations reinforce the sensitivity of pollen to thermal stress and provide a physiological basis for the biochemical and metabolomic differences observed in this study. Pollen responses to environmental stress are known to manifest at multiple molecular levels, including transcriptome, proteome, and metabolome alterations (Chaturvedi et al., 2021), underscoring the complexity of pollen stress adaptation. Consistent with this, our results suggest that released pollen remains metabolically and biochemically active under temperature stress, with stress-dependent modulation of cellular processes.

Under heat stress, pollen typically attempts to maintain cellular homeostasis through adjustments in protein conformation, membrane fluidity, cytoskeletal organization, and redox balance, all of which are critical for sustaining metabolic stability (Zinn et al.,2010). In our study, reduced esterase activity under heat stress, a key biochemical indicator of pollen viability, is indicative of compromised membrane integrity and cellular function, consistent with previous observations in *Oryza sativa* and *Arabidopsis thaliana* (Endo et al., 2009). A major consequence of heat stress is excessive ROS accumulation, which can damage lipids, proteins, and nucleic acids, and ultimately impair pollen germination and viability. Similarly, increased ROS levels under heat stress have been reported in *Triticum aestivum* pollen, supporting our observation that oxidative stress is a common consequence of elevated temperatures in reproductive tissues (Sarkar et al., 2021)._In line with this, our previous work showed that unreleased pollen exhibited minimal ROS accumulation, whereas released pollen displayed elevated ROS levels, suggesting that environmental exposure during dispersal acts as a trigger for ROS generation through multiple stress-responsive signaling pathways (Rutuparna et al., 2022). Together, these findings indicate that oxidative stress is closely associated with reduced pollen viability under heat stress, particularly in released pollen directly exposed to adverse environmental conditions.

A marked difference in proline accumulation was observed between cold– and heat-stressed pollen, with significantly higher proline levels under cold stress compared to heat stress. This stress-dependent accumulation pattern aligns with the well-established role of proline as a protective metabolite contributing to pollen viability by stabilizing cellular membranes, maintaining osmotic balance, and preventing protein denaturation during low-temperature exposure (Mattioli et al., 2018; Kiran et al., 2021; Goel et al., 2023). Beyond its osmoprotective function, proline also serves as a readily mobilizable energy source, supporting pollen resilience during dehydration and temperature fluctuations, as reported previously (Mattioli et al., 2018).

In contrast, the reduced proline content observed under heat stress suggests perturbation of proline homeostasis, potentially arising from impaired biosynthesis or reduced transporter-mediated uptake. Supporting this interpretation, studies in tomato anthers have shown that prolonged exposure to elevated temperatures represses proline transporter activity during meiosis, thereby limiting proline accumulation and negatively affecting pollen development and fertility (Firon et al., 2012; Chaturvedi et al., 2021). Similar temperature-dependent regulation of proline metabolism has been documented in barley and other cereals, where heat stress induces downregulation of genes involved in proline biosynthesis (Zenda et al., 2022). These findings suggest that enhanced proline accumulation under cold stress is associated with improved pollen survival, whereas proline depletion under heat stress may compromise membrane integrity and reduce pollen viability, highlighting the importance of tightly regulated proline metabolism for reproductive success under temperature extremes.

Phytohormones play a critical role in regulating pollen development and fertility, particularly under environmental stress conditions (Bhatt et al., 2020). To examine their contribution to pollen stress adaptation, we quantified eight phytohormones – abscisic acid (ABA), indole-3-acetic acid (IAA), indole-3-butyric acid (IBA), 1-naphthaleneacetic acid (NAA), gibberellic acid (GA□), kinetin, salicylic acid (SA), and methyl jasmonate – under cold (R15) and heat (R35) stress conditions. All analyzed hormones exhibited measurable, temperature-dependent variation, indicating a complex hormonal cross-talk underlying pollen stress responses (Hirano et al., 2008). Auxins such as IAA, IBA, and NAA are essential regulators of pollen development and reproductive competence (Liu et al., 2024). In the present study, IAA levels were higher in heat-stressed pollen (0.722 nmol g ¹) than in cold-stressed pollen (0.476 nmol g ¹), suggesting enhanced auxin-associated signalling responses under elevated temperatures, consistent with reports of heat-induced auxin modulation in reproductive tissues (Liu et al., 2024; Wu et al., 2025).Similarly, IBA accumulation increased under heat stress, whereas NAA levels were comparatively lower under cold stress, suggesting temperature-dependent regulation of auxin metabolism. Among the analyzed phytohormones, gibberellic acid (GA_3_) and salicylic acid (SA) exhibited the most pronounced changes. GA_3_ was the most abundant hormone overall, with concentrations of 479.86 nmol g ¹ under cold stress (R15) and 358.2 nmol g ¹ under heat stress (R35), indicating a potential role for gibberelins in maintaining pollen function under low-temperature conditions (Chaturvedi et al., 2021). The observed reduction in GA levels under heat stress further highlights its temperature sensitivity, in agreement with studies reporting GA repression under abiotic stress through altered auxin–ABA balance (Kaur et al., 2021). Salicylic acid, a key regulator of plant defense and abiotic stress tolerance, was also elevated in heat-stressed pollen, indicating its involvement in maintaining pollen viability under high-temperature exposure (Chaturvedi et al., 2021). ABA levels also increased under heat stress (0.047 nmol g ¹) compared to cold stress (0.028 nmol g ¹), consistent with its established role in mediating stress tolerance and regulating ROS-related signalling (Hsu et al., 2021).

In contrast, methyl jasmonate showed no significant difference between cold (0.009 nmol g ¹) and heat (0.008 nmol g ¹) stress conditions, aligning with previous reports that associate its primary function with anther dehiscence rather than direct temperature stress responses in *Arabidopsis thaliana* (Peng et al., 2013). Overall, these findings highlight that different phytohormones contribute distinctively to pollen thermotolerance, with GA and SA potentially supporting pollen survival under cold and heat stress respectively, while auxins and ABA exhibit temperature-dependent adjustments that fine-tune pollen metabolism and fertility in response to fluctuating environmental conditions.

We observed distinct, temperature-dependent variations in polyamine levels under cold and heat stress, with putrescine being the most abundant polyamine among the four analyzed. Putrescine is known to play a crucial role in maintaining pollen integrity and viability, thereby supporting successful fertilization and pollen tube growth (Aloisi et al., 2016). In the present study, putrescine levels were consistently higher than those of other polyamines under both cold and heat stress, suggesting its importance in sustaining pollen metabolism during stress. This observation aligns with previous reports showing that putrescine accumulation often coincides with reduced levels of spermine and spermidine, under heat stress conditions (Upadhyay et al., 2020). Cadaverine content showed a reduction under heat stress compared with cold stress, suggesting that its contribution to pollen stress adaptation may be limited under temperature extremes, as reported previously (Jancewicz et al., 2016). Although putrescine was elevated under both temperature treatments, spermidine showed a modest increase under heat stress, suggesting a possible role in supporting pollen viability under moderately elevated temperatures (Santiago and Sharkey, 2019) In contrast, spermine levels declined under both cold and heat stress, suggesting that temperature exposure perturbs polyamine homeostasis in released pollen. Collectively, these findings indicate that differential regulation of polyamine pools occurs in response to temperature stress, with elevated levels of putrescine and spermidine potentially contributing to the maintenance of pollen viability, while reductions in spermine and cadaverine may reflect stress-induced shifts in polyamine metabolism. Such rebalancing of polyamine composition is likely an important component of pollen stress adaptation, helping to preserve cellular integrity and function under adverse temperature conditions (Alcazar et al., 2010).

Given the pronounced biochemical changes observed in sugars, phytohormones, and polyamines, we next performed untargeted metabolomic profiling to obtain a broader and integrated understanding of the metabolic adjustments supporting pollen survival under cold and heat stress. This systems-level approach complements the targeted biochemical analyses by capturing coordinated shifts in primary and secondary metabolism that underlie pollen viability and stress tolerance. In particular, untargeted metabolomic investigations in wind-pollinated pollen under abiotic stress have rarely been reported. Cold stress is known to markedly alter the physiological and metabolic processes of plants, particularly those associated with pollen viability and fertility (Müller and Rieu, 2016).

Among the most prominently upregulated metabolites in cold-stressed pollen were berberine, palmitic acid, and glycocholic acid, each contributing uniquely to cellular stress adaptation. The accumulation of berberine, a bioactive alkaloid with antioxidant properties, is consistent with the higher pollen viability observed under cold stress and suggests activation of secondary metabolic defenses that mitigate oxidative damage, as reported in other stressed reproductive tissues (Singh et al., 2021). Although berberine is not traditionally considered a pollen-specific compound, its induction under cold stress likely reflects a broader stress protection strategy mediated by secondary metabolism (Liu et al., 2024). This observation aligns with the comparatively higher pollen viability and lower oxidative damage observed under cold stress in our viability assays.

The upregulation of palmitic acid, a saturated fatty acid, suggests enhanced membrane remodelling, a canonical response that preserves membrane stability and fluidity under temperature extremes (Huby et al., 2020). This lipid remodeling is consistent with the higher pollen viability observed under cold stress and supports the notion that membrane stabilization is a key determinant of pollen survival. Increased palmitic acid levels have also been shown to be associated with improved cold stress resilience through the modulation of membrane lipid saturation (Tian et al., 2022). Other upregulated metabolites, including diallyl trisulfide and hydrangenoside C, are implicated in plant defense signaling and oxidative stress management, corroborating their functional enhancement under chilling, as shown in several crop systems (Whitehead and bowers, 2013). The accumulation of dihydrosphingosine, a sphingolipid precursor, essential for membrane integrity, further supports enhanced protection against oxidative and mechanical stress in cold-stressed pollen, consistent with studies on pollen and anther resilience to stress (Esposito et al., 2019). Similarly, the increased abundance of 7-hydroxy-4-methylcoumarin, an antioxidant phenolic compound, suggests improved ROS scavenging capacity, which may contribute to maintaining pollen viability under cold conditions (Vianna et al., 2012). Elevated sitosterol under cold stress further indicates membrane reorganization and signaling adjustments, required for stress adaptation (Vogel et al., 2025). Osajin, and ormosanine, derived from flavonoid and alkaloid biosynthetic pathways, likely represent additional layers of biochemical defense, consistent with the well-established roles of phenolic compounds and alkaloids in plant protective metabolism (Dixon and Paiva, 1995; Wink, 2010; Agati et al., 2012;). Kynurenine, a regulator of auxin-related pathways, may facilitate the hormonal adjustments observed in cold-stressed pollen, linking metabolomic changes with the hormone profiles quantified in this study (Suzuki et al., 2015). Collectively, the predominance of defense-related, antioxidant, and membrane-stabilizing metabolites under cold stress reflects a coordinated metabolic reprogramming that supports pollen viability, aligns with the higher proline accumulation and favorable hormonal balance observed under cold conditions, and ultimately promotes successful fertilization under low-temperature environments (Huang et al., 2022).

The downregulated metabolites collectively reflect a complementary metabolic adjustment in response to temperature stress. The marked reduction of isopalmitic acid suggests suppressed lipid biosynthesis and remodeling activity, which may correlate with lower reactive oxygen species (ROS) production in cold-stressed pollen relative to heat-stressed pollen, indicating a strategic modulation of oxidative stress(Zhao et al., 2018). This is consistent with our biochemical observations showing reduced oxidative damage under cold stress compared to heat stress. Consequently, a reduced accumulation of antioxidant flavonoids may be sufficient under cold conditions in this system, potentially reflecting a lower oxidative burden or compensation by alternative protective mechanisms, such as enhanced osmolyte accumulation and membrane stabilization. This is consistent with observed decreases in metabolites such as karanjin, osthol, and DAH-3-keto-4-en, which are typically involved in flavonoid biosynthesis and pigmentation pathways (Agati et al., 2012). Similarly, the downregulation of alizarin, an anthraquinone known for its pigment-related defense, indicates a reduced reliance on pigmentation-based stress responses during cold exposure, consistent with observations from cold stress metabolomic studies in other plant systems (Langa et al., 2021). The reduced abundance of reticulol and carnosine further suggests diminished oxidative stress signaling or lower antioxidant demand in cold-stressed pollen, aligning with earlier findings in wheat and tomato pollen subjected to temperature stress (Sharma and Nayyar, 2016).

The combined multivariate and univariate analyses revealed a distinct metabolic signature associated with cold– and heat-stress responses in pollen. Sparse partial least squares–discriminant analysis (sPLS-DA) identified a subset of metabolites with strong discriminatory power between temperature conditions, many of which overlapped with features selected by both differential expression analysis and PLS-DA (VIP > 1.2), highlighting the consistency and robustness of this metabolic marker panel. This high-confidence subset included palmitic acid, dihydrosphingosine, and cupressuflavone primarily associated with cold stress, as well as karanjin, 2′-deoxyadenosine, biliverdin, 3′,4′,5,7-tetrahydroxyflavone, piperidine, caryophyllene, and theobromine, which are predominantly linked to heat-stress responses. The directionality of sPLS-DA loadings further supported stress-specific metabolic patterns, with negative loaded metabolites (e.g., palmitic acid and dihydrosphingosine) enriched under cold stress, and positively loaded metabolites (e.g., theobromine and caryophyllene) associated with heat stress. These trends were consistent with fold-change-based differential expression results, validating the biological relevance of the selected metabolites. While the biochemical roles of many metabolites have been discussed individually, this integrative multivariate perspective highlights a coordinated metabolic reprogramming that supports pollen adaptation to contrasting temperature stresses. Notably, a few heat-stress-enriched metabolites, such as karanjin, 2′-deoxyadenosine, and biliverdin emerged as promising candidates for further investigation into their potential roles in pollen heat perception, signalling and defense mechanisms.

The enriched metabolic pathways identified from the pathway analysis highlight key biochemical and regulatory mechanisms underlying pollen responses to cold and heat stress, providing pathway-level support for the metabolite-level changes observed in this study. Purine metabolism is significantly enriched, indicating elevated nucleotide turnover and recycling. Such processes are essential for maintaining cellular energy balance and the biosynthesis of stress-related signalling molecules, such as abscisic acid, which play a critical role in pollen activation and stress adaptation under temperature extremes (Chen et al., 2024; Rezaul et al., 2019). Enhanced purine flux under heat stress may facilitate increased energy demand associated with stress responses, including the synthesis of heat shock proteins and osmoprotectants such as sucrose and raffinose. Conversely, rapid nucleotide recycling under cold stress likely supports energy and nitrogen demand associated with pollen maintenance and activation, consistent with previous observations (Chen et al., 2024). Arginine biosynthesis and arginine-proline metabolism were also prominently enriched, connecting pathway-level changes to the elevated proline accumulation and altered polyamine profiles observed under cold stress. These pathways play central roles in nitrogen metabolism and osmoprotection, linking proline accumulation and polyamine biosynthesis, which is essential for membrane stabilization and ROS scavenging under temperature stress conditions (Szabados and Savoure, 2010; Alcazar et al., 2010). Enrichment of glutathione metabolism further highlights the importance of redox homeostasis and ROS detoxification mechanisms in protecting pollen against oxidative damage induced by heat and cold stress (Noctor et al., 2012; Foyer and Noctor, 2011). Similarly, enrichment of vitamin B6 (pyridoxine) metabolism suggests a role in antioxidant defense and the regulation of enzymatic processes associated with pollen viability under stress conditions (Titiz et al., 2006).

The glyoxylate and dicarboxylate metabolism pathway is closely linked to central carbon metabolism and photorespiration, providing metabolic flexibility that supports cellular energy balance under stress conditions (Wingler et al., 2000; Jiang et al., 2023). Enrichment of beta-alanine metabolism contributes to coenzyme A biosynthesis and associated metabolic buffering, supporting cellular survival and antioxidant capacity under temperature extremes (Parthasarathy et al., 2019; Wu et al., 2025). Tyrosine-derived pigments such as betalains are well-recognized antioxidants that function as direct ROS scavengers and contribute to cellular redox balance during temperature stress (Li et al., 2019). Enrichment of the citrate cycle (TCA cycle) reflects increased respiratory activity and energy production required to sustain stress responses (Xu and Fu, 2022). Zeatin biosynthesis enrichment further suggests a role of cytokinins in coordinating growth-defense trade-offs during pollen stress adaptation, orchestrating hormonal regulation along with auxins, ABA, and gibberellins to fine-tune hormonal regulation under fluctuating temperature conditions (Zwack and Rashotte, 2015; Pavlu et al., 2018; Chaturvedi et al., 2021). Overall, these pathways show that pollen adapts to temperature stress by regulating membrane stability, antioxidant defenses, hormone signaling, and energy metabolism to maintain viability and reproductive success under adverse temperature conditions.

The agreement between univariate analyses (differential expression analysis), multivariate approaches (PLS-DA VIP scores and sPLS-DA) and pathway-level enrichment strengthens confidence in the biological relevance of the identified metabolic changes. However, this study has certain limitations, including a relatively small sample size and the semi-quantitative nature of untargeted metabolomics, which may limit the detection of low-abundance or transient stress-responsive metabolites. In addition, species-specific metabolic traits and environmental variability may influence pollen metabolic profiles and should be considered when extrapolating these findings.

Despite these constraints, the integrative analytical framework employed here enabled the identification of a focused subset of metabolites with strong discriminatory potential between cold– and heat-stress conditions. These metabolites provide insight into the coordinated metabolic reprogramming that supports pollen viability under adverse thermal environments. Future studies incorporating larger sample sizes, independent biological validation, and multi-omics integration (including transcriptomic and proteomic analyses), along with expanded sampling across species, and field conditions, will be required to rigorously evaluate the reproducibility, biological relevance, and potential utility of these metabolites as robust stress indicators.

## Conclusions

This study provides insights into how pollen metabolism is differentially modulated under cold and heat stress during the critical phase of pollen release and dispersal. Our findings improve current understanding of pollen physiology, metabolism, homeostasis, and adaptive responses to temperature extremes associated with wind pollination. We demonstrate that released pollen retains partial metabolic activity and undergoes specific biochemical and metabolic adjustments that contribute to stress resilience. Integrative biochemical and untargeted metabolomic profiling revealed temperature-dependent alterations in key metabolites and metabolic pathways associated with pollen tolerance to cold and heat stress. Collectively, these results highlight distinct molecular and biochemical strategies employed by pollen to mitigate temperature-induced damage and maintain viability and reproductive function. This work provides a foundation for future studies aimed at validating stress-responsive metabolic indicators and identifying metabolic targets to improve pollen performance and crop fertility under changing climatic conditions.

## Acknowledgements

Jena Rutuparna and Irfan Ahmad Ghazi thank the Department of Science and Technology (DST), Government of India, for the award of DST-INSPIRE-JRF and financial support through a DST-SERB project, respectively. The authors gratefully acknowledge the scientific assistance and guidance of the late Prof. K. Seshagirirao. The authors also thank the UoH Metabolomics Facility, School of Life Sciences, University of Hyderabad, for technical support.

## Author contributions

JR conceived, designed, and performed the experiments and analyzed the results. IAG designed the experiments and contributed materials and reagents. JI performed mass spectrometry, acquired the metabolomic data, and conducted data analysis. AA performed experiments and analyzed the data. JR, JI, AA, Divya, and Raj wrote the manuscript. IAG reviewed and edited the manuscript. All authors read and approved the final manuscript.

## Use of Artificial Intelligence in the writing process

ChatGPT 5.2 (OpenAI) was used solely for language editing, including improvements to grammar, clarity, and readability of the manuscript. The tool was not used to generate scientific content, interpret data, draw conclusions, or create figures. All scientific ideas, analyses, results, and interpretations presented in this manuscript are the sole responsibility of the authors, who reviewed and verified the final content.

