## Supplementary Figure for "Comparative metabolomics of released pollen during dispersal reveals metabolic adaptations to cold and heat stress"

*Corresponding author: Jena Rutuparna

**Co-corresponding author: Irfan Ahmad Ghazi

**Supplementary Figure S1** HPLC-based quantification of sugars in pollen under cold (R15) and heat (R35) stress conditions. Glucose, sucrose, and fructose concentrations are expressed as mg g⁻¹ fresh weight.

**Supplementary Figure S2** Proline estimation in pollen under temperature stress.
**Supplementary Figure S3** Polyamine profiling in pollen under cold-stressed pollen (CSPP; R15) and heat-stressed pollen (HSPP; R35) conditions.

**Supplementary Figure S4** Principal Component Analysis (PCA) of metabolomic profiles

**Supplementary Figure S5** Hierarchical clustering heatmap of top 50 significantly altered metabolites across cold- and heat-stressed pollen samples.

**Supplementary Figure S6** PLS-DA analysis

**Supplementary Figure S7** sPLS-DA analysis

**Supplementary Figure S8** Pathway enrichment network showing functionally grouped and significantly enriched metabolic pathways based on shared metabolites or functional similarity

Node size reflects pathway impact; node colour represents statistical significance (darker = more significant); edges indicate shared metabolites between pathways

**Supplementary Table S1** Top metabolites ranked by Variable Importance in Projection (VIP) scores from PLS-DA analysis. VIP scores reflect the contribution of each metabolite to group separation. Metabolites with VIP > 1 are generally considered significant contributors.

**Supplementary Figure S1** HPLC-based quantification of sugars in pollen under cold (R15) and heat (R35) stress conditions. Glucose, sucrose, and fructose concentrations are expressed as mg g⁻¹ fresh weight.

Sugars content (mg/g)

|  | CSPP | HSPP |
| --- | --- | --- |
| Glucose | 9.705 | 6.78 |
| Sucrose | 10.133 | 10.932 |
| Fructose | 0.587346 | 0.2357 |

**Supplementary Figure S2** Proline estimation in pollen under temperature stress.

A

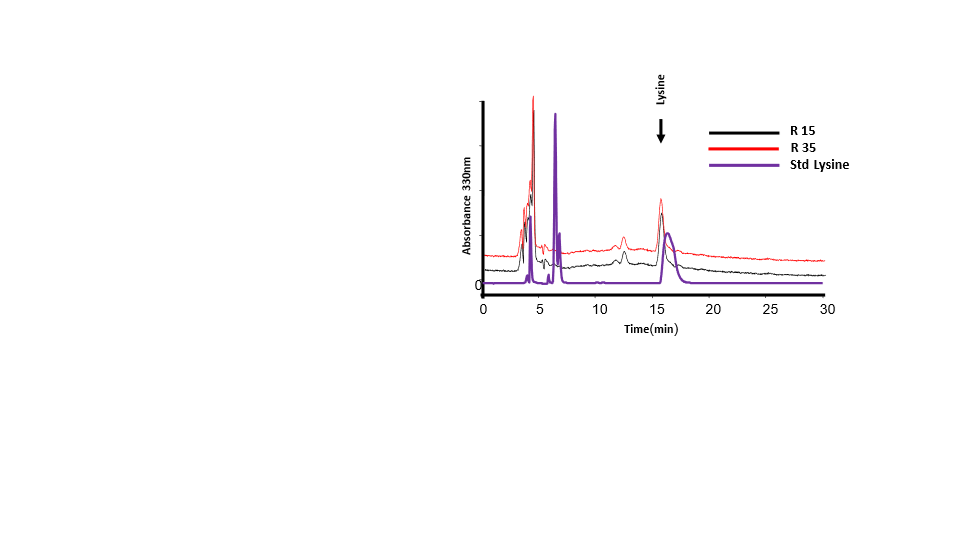

B

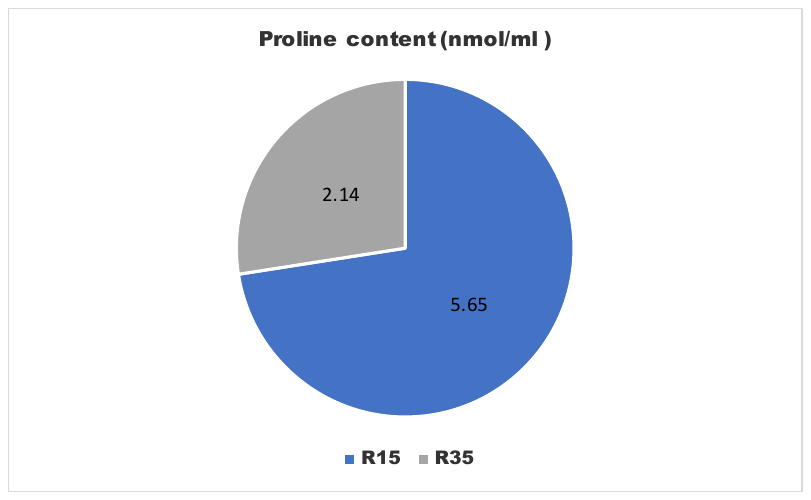

(A) Representative HPLC chromatogram for amino acid analysis.
(B) Proline quantification in pollen exposed to cold (R15) and heat (R35) stress, expressed as nmol mL⁻¹.

**Supplementary Figure S3** Polyamine profiling in pollen under cold-stressed pollen (CSPP; R15) and heat-stressed pollen (HSPP; R35) conditions

A

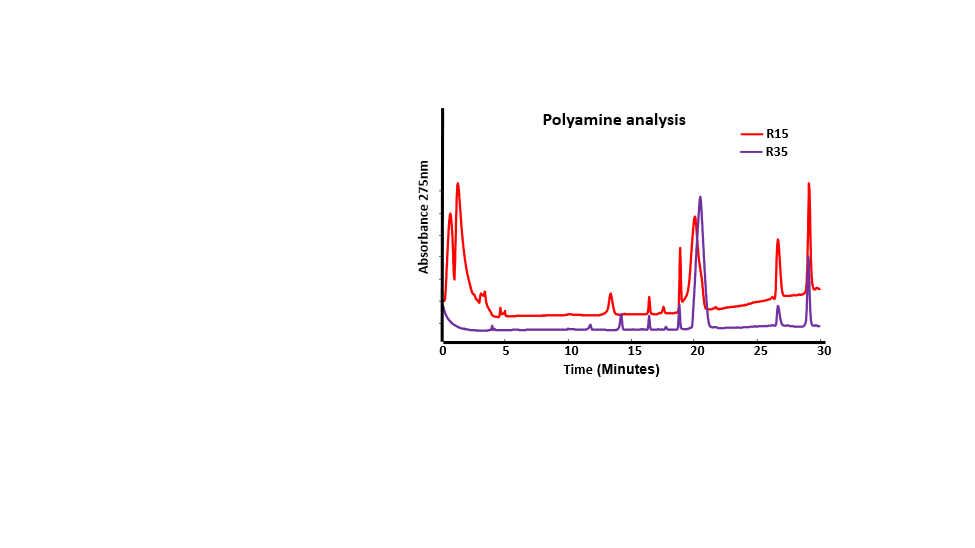

**B**

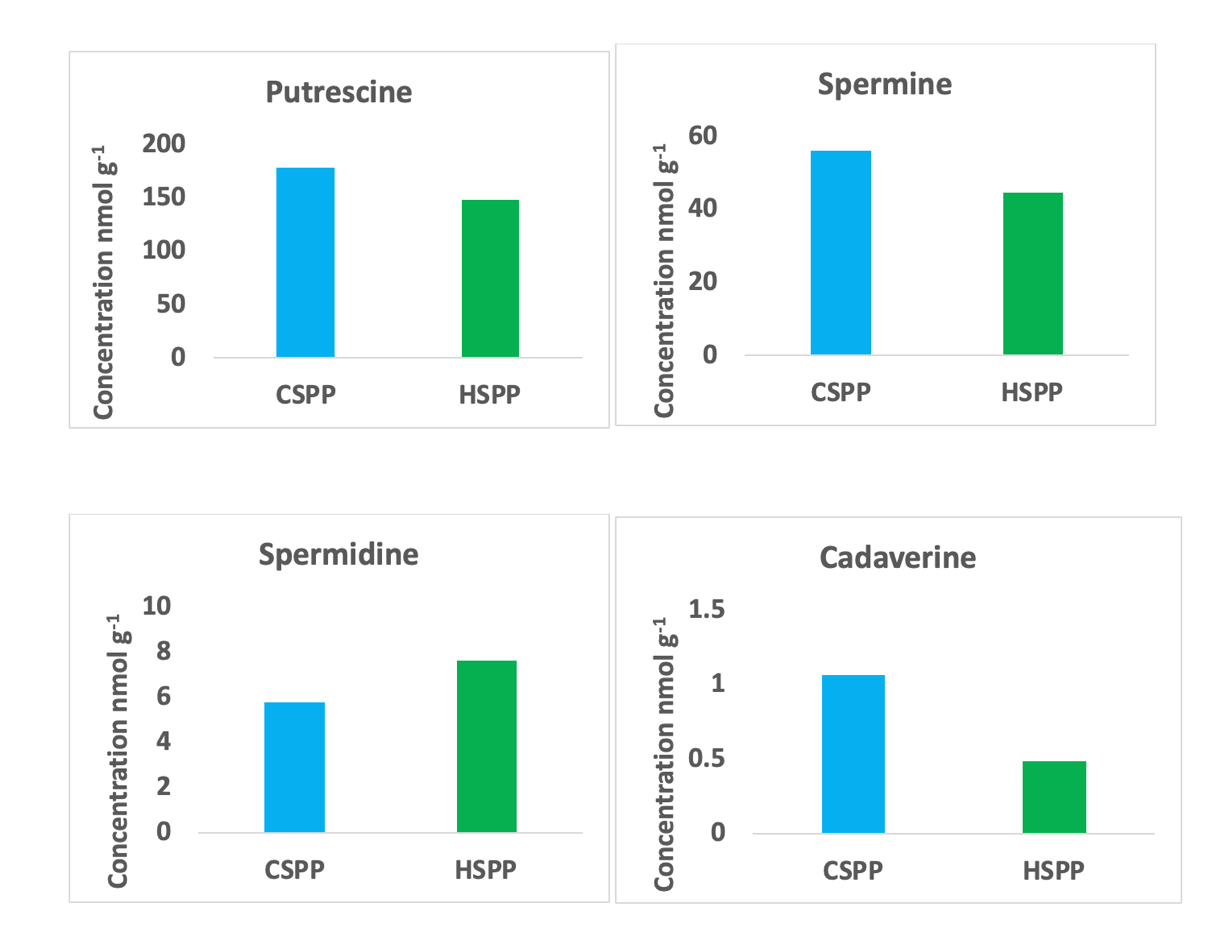

(A) Representative HPLC chromatogram for polyamine analysis

(B) Quantification results for Putrescine, Spermine, Spermidine and Cadaverine

**Supplementary Figure S4** Principal Component Analysis (PCA) of metabolomic profiles

A

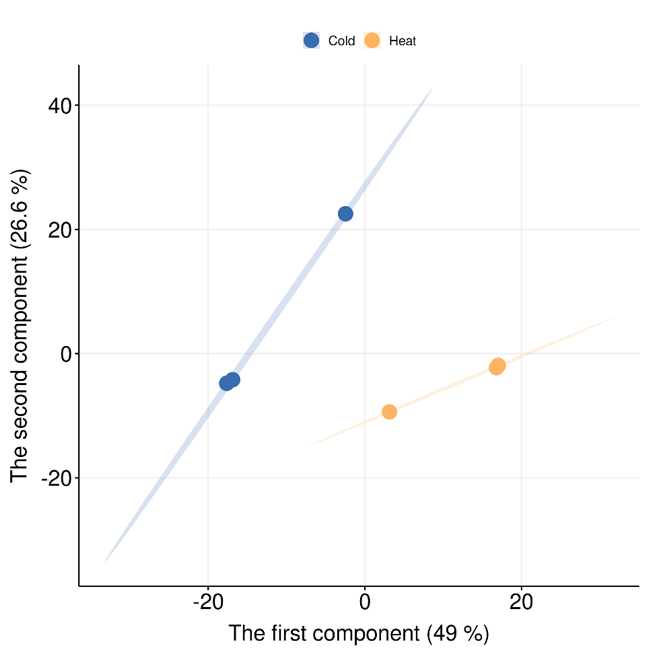

B

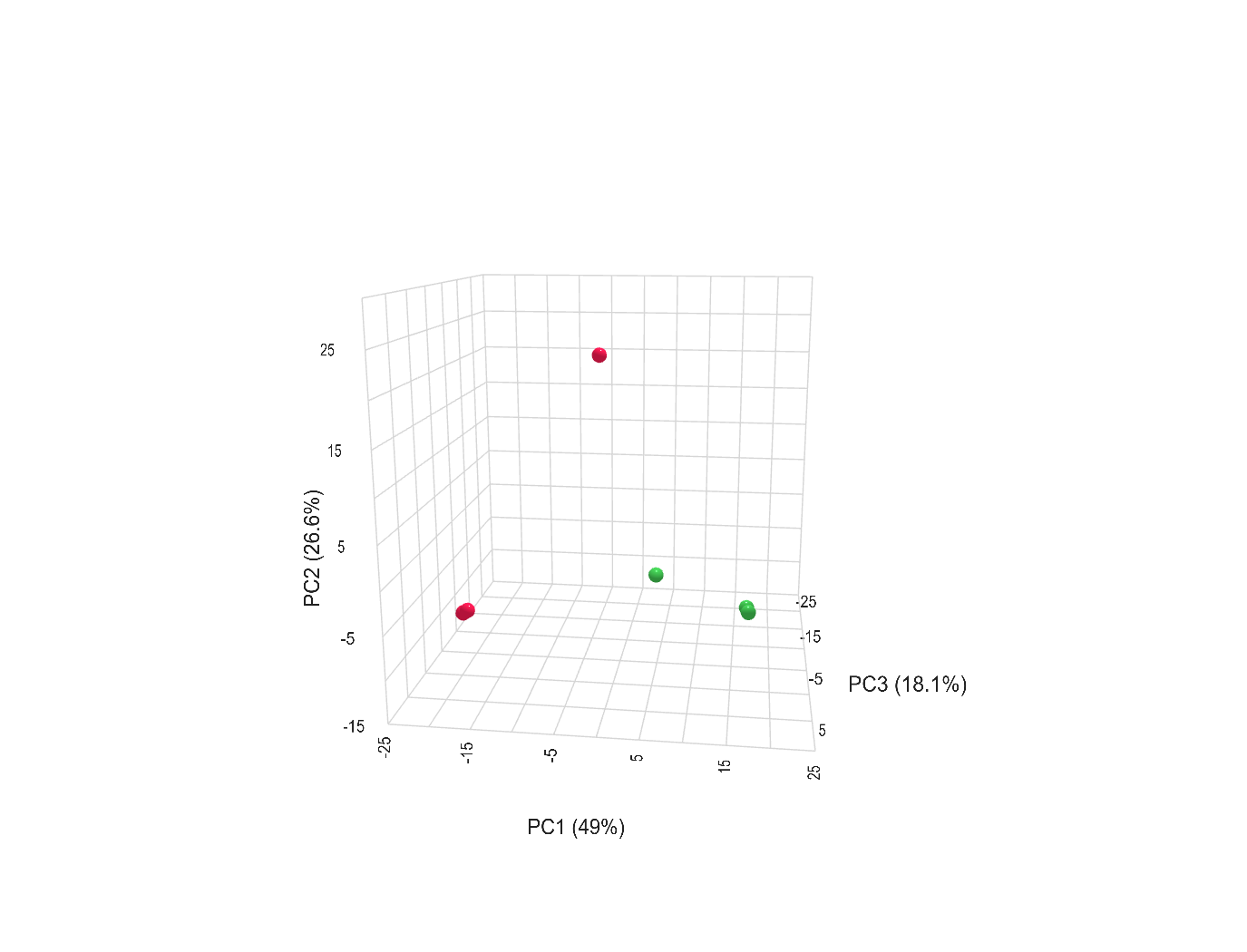

(A) 2D PCA score plot and
(B) 3D PCA plot showing separation between cold- and heat-stressed pollen samples based on all detected metabolites. Each point represents a biological replicate

**Supplementary Figure S5** Hierarchical clustering heatmap of top 50 significantly altered metabolites across cold- and heat-stressed pollen samples.

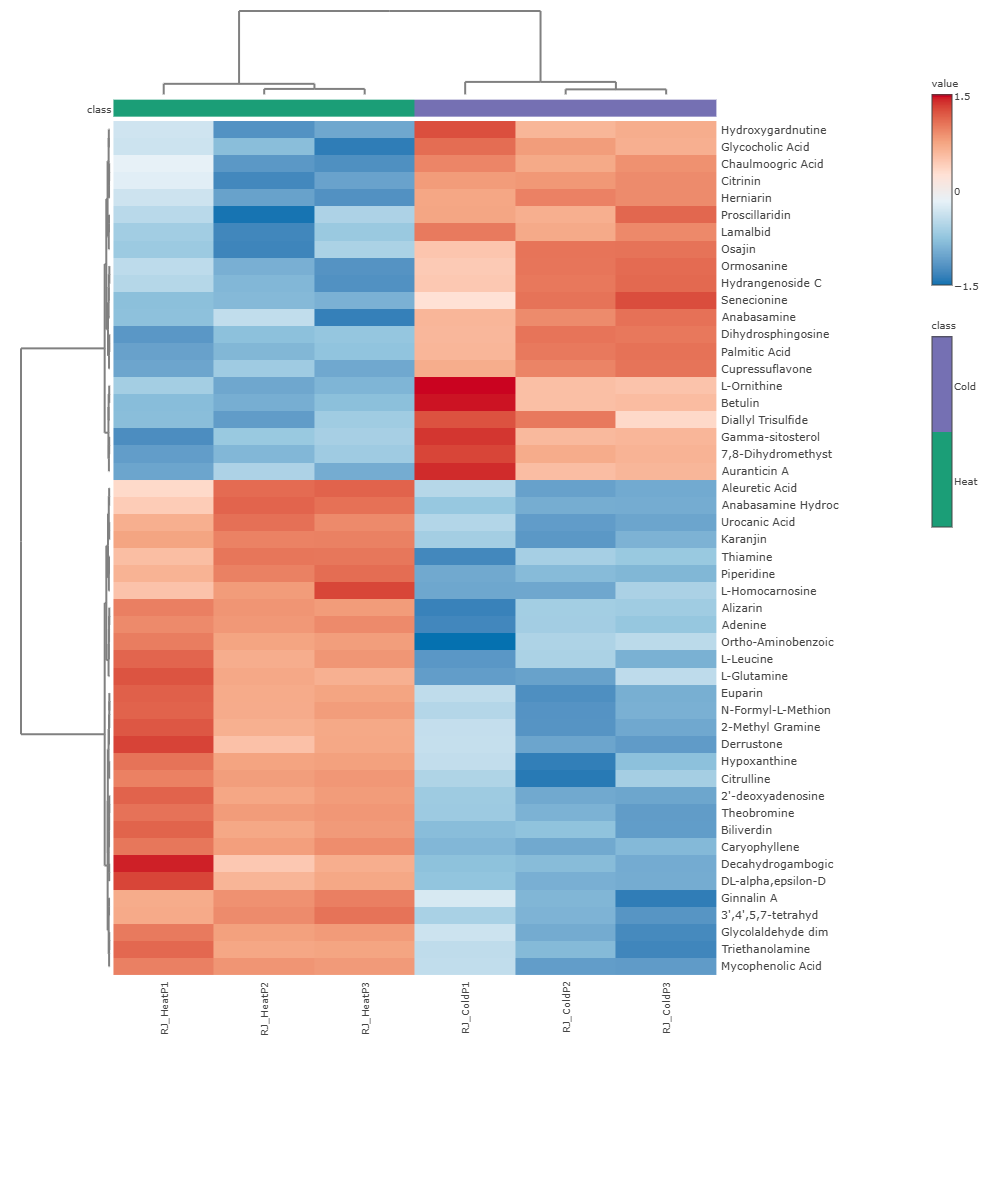

**Supplementary Figure S6.** Partial least squares–discriminant analysis (PLS-DA).

A

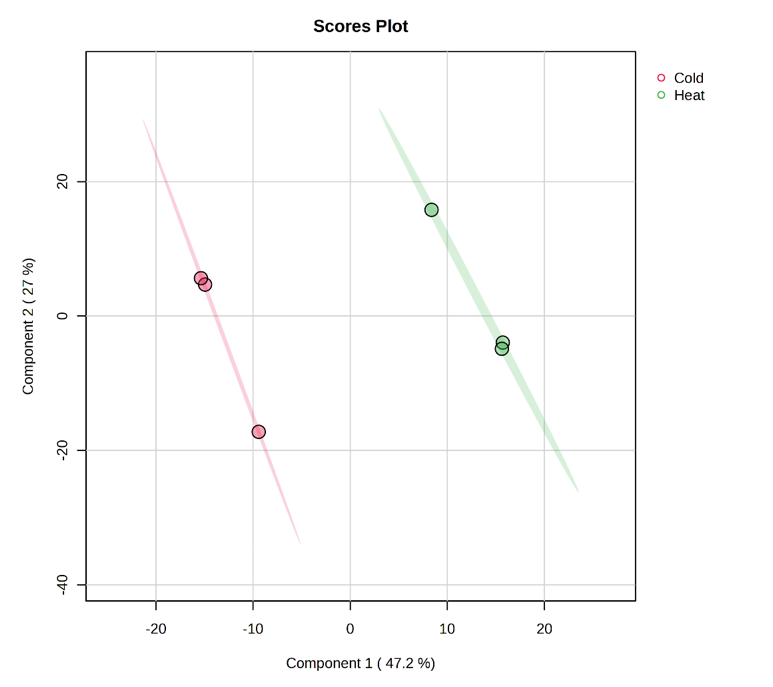

B

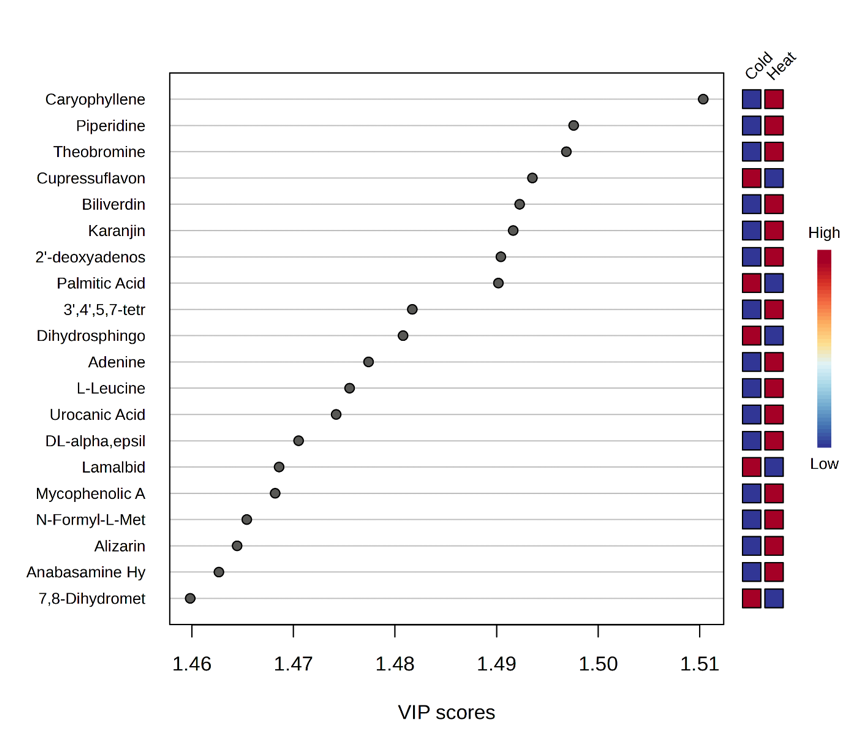

C

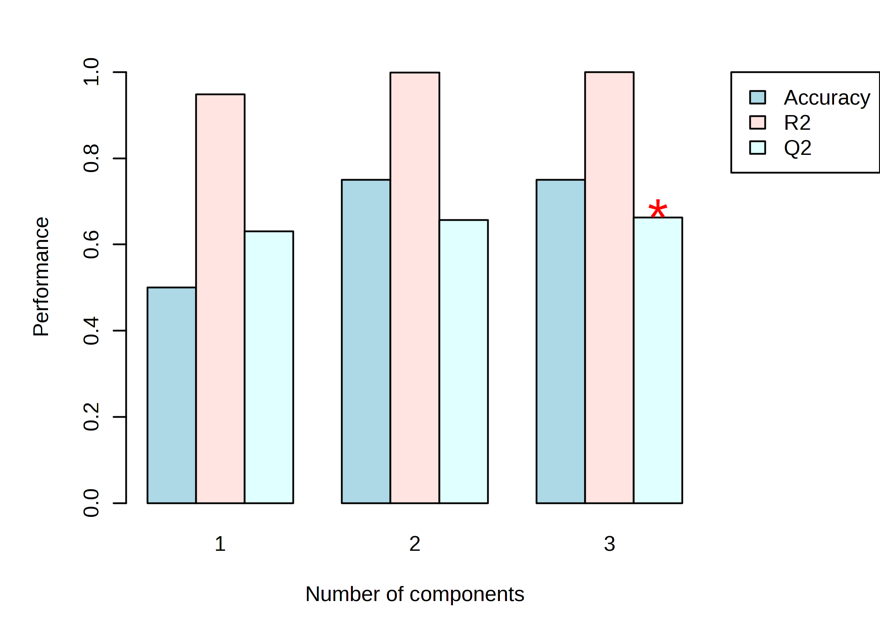

PLS-DA cross-validation details

| **Measure** | **1 component** | **2 components** | **3 components** |
| --- | --- | --- | --- |
| Accuracy | 0.5 | 0.75 | 0.75 |
| R2 | 0.9482 | 0.99853 | 0.99994 |
| Q2 | 0.63034 | 0.65686 | 0.66266 |

(A) Scores plot showing group separation between cold- and heat-stressed pollen samples
(B) Variable Importance in Projection (VIP) plot indicating top contributing metabolites
(C) Cross-validation results displaying model performance (R² and Q²)

**Supplementary Figure S7.** Sparse partial least squares–discriminant analysis (sPLS-DA).

A

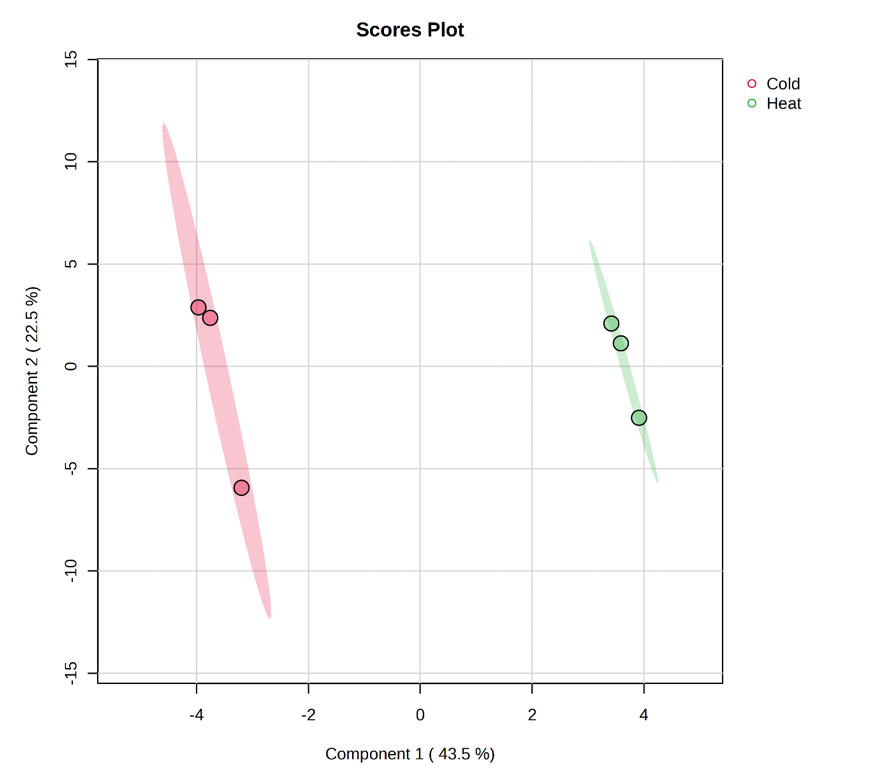

B

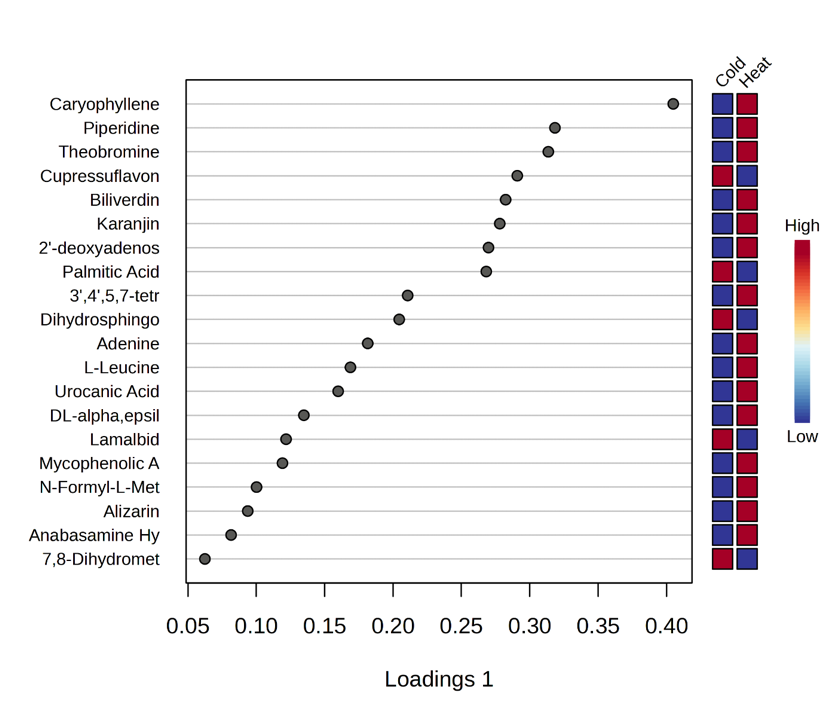

C

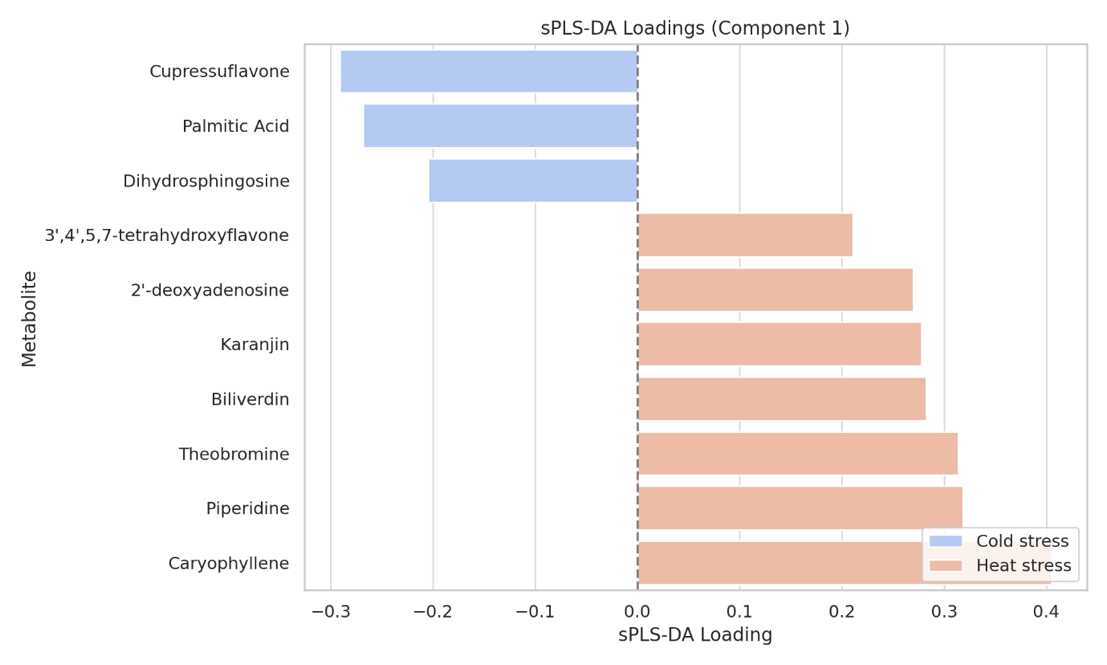

(A) Scores plot showing group separation
(B) Bar plot of key discriminative metabolites selected in Component 1 and Component 2
(C) Loadings of the top ten metabolites contributing to group discrimination

**Supplementary Figure S8.** Pathway enrichment network of significantly enriched metabolic pathways.

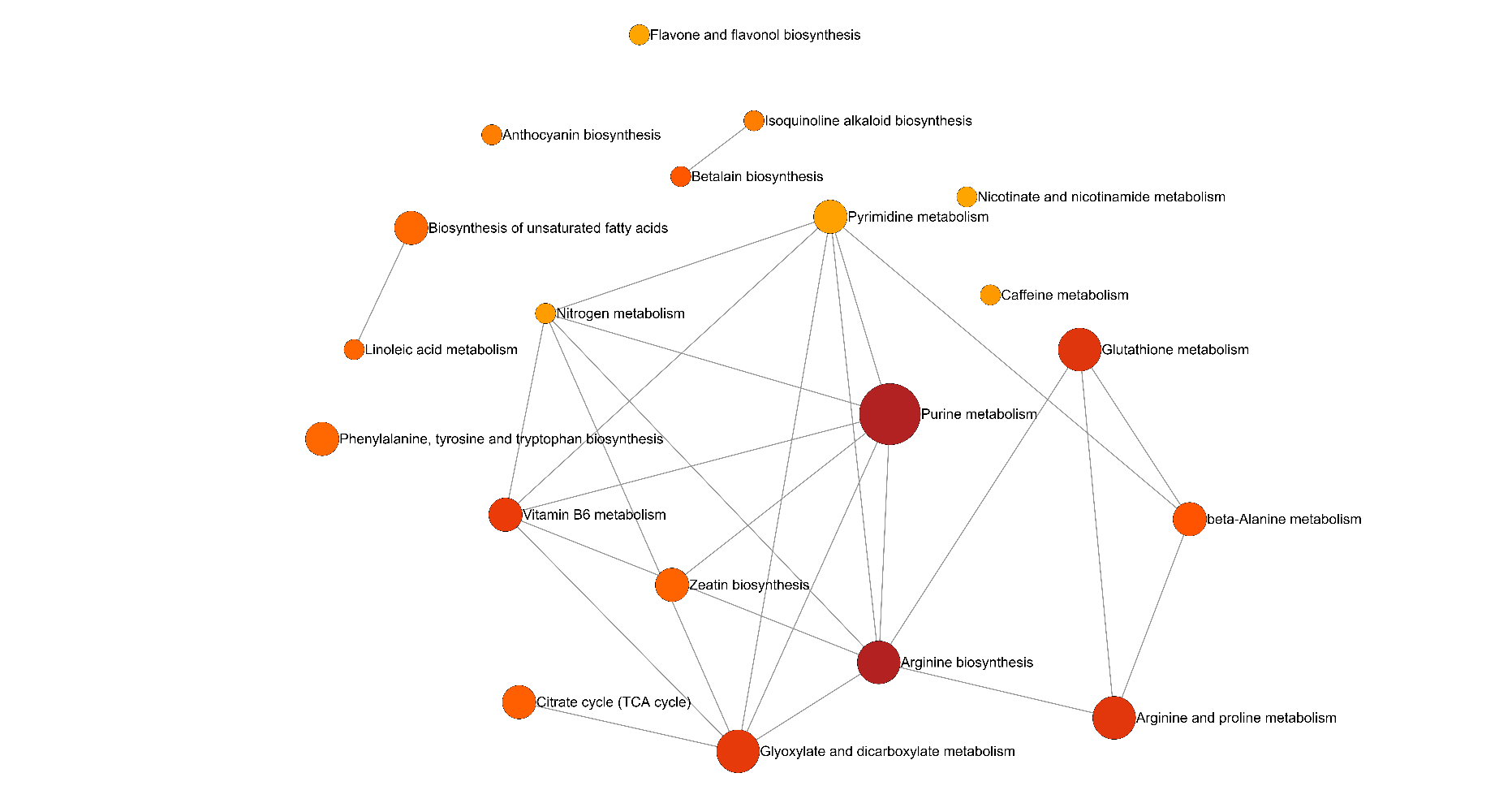

Node size reflects pathway impact; node colour represents statistical significance (darker colours indicate lower p-values); edges indicate shared metabolites between pathways.

**Supplementary Table 1.** Top metabolites ranked by Variable Importance in Projection (VIP) scores from PLS-DA analysis. VIP scores reflect the contribution of each metabolite to group separation. Metabolites with VIP > 1 are generally considered significant contributors.

| Metabolite | Component 1 | Component 2 |
| --- | --- | --- |
| Caryophyllene | 1.5103 | 1.4767 |
| Piperidine | 1.4976 | 1.4608 |
| Theobromine | 1.4969 | 1.4618 |
| Cupressuflavone | 1.4935 | 1.4583 |
| Biliverdin | 1.4923 | 1.4602 |
| Karanjin | 1.4916 | 1.454 |
| 2'-deoxyadenosine | 1.4904 | 1.4568 |
| Palmitic Acid | 1.4902 | 1.4547 |
| 3',4',5,7-tetrahydroxyflavone | 1.4817 | 1.444 |
| Dihydrosphingosine | 1.4808 | 1.4467 |
